# The optimal odor-receptor interaction network is sparse in olfactory systems: Compressed sensing by nonlinear neurons with a finite dynamic range

**DOI:** 10.1101/464875

**Authors:** Shanshan Qin, Qianyi Li, Chao Tang, Yuhai Tu

## Abstract

There are numerous different odorant molecules in nature but only a relatively small number of olfactory receptor neurons (ORNs) in brains. This “compressed sensing” challenge is compounded by the constraint that ORNs are nonlinear sensors with a finite dynamic range. Here, we investigate possible optimal olfactory coding strategies by maximizing mutual information between odor mixtures and ORNs’ responses with respect to the bipartite odor-receptor interaction network (ORIN) characterized by sensitivities between all odorant-ORN pairs. We find that the optimal ORIN is sparse – a finite fraction of sensitives are zero, and the nonzero sensitivities follow a broad distribution that depends on the odor statistics. We show that the optimal ORIN enhances performances of downstream learning tasks (reconstruction and classification). For ORNs with a finite basal activity, we find that having a basal-activity-dependent fraction of inhibitory odor-receptor interactions increases the coding capacity. All our theoretical findings are consistent with existing experiments and predictions are made to further test our theory. The optimal coding model provides a unifying framework to understand the peripheral olfactory systems across different organisms.

## I. INTRODUCTION

Animals rely on their olfactory systems to detect, discriminate and interpret external odor stimuli to guide their behavior. Natural odors are typically mixtures of different odorant molecules whose concentrations can vary over several orders of magnitude [1–3]. Remarkably, animals can distinguish a large number of odorants and their mixtures by using a relatively small number of odor receptors (ORs) [4, 5]. For example, humans have only ~ 300 ORs [6, 7], and the often cited number of odors that can be distinguished is ~ 10000 [8], the real number may be even larger [9] (see also [10] and [11]). Humans can also distinguish odor mixtures with up to 30 different compounds [12]. In comparison, the highly olfactory lifestyle and exquisite olfactory learning ability of the fly is afforded by only ~ 50 ORs [4, 13]. The olfactory system achieves such remarkable ability through a combinatorial code in which each odorant is sensed by multiple receptors and each receptor can be activated by many odorants [14–16]. In both mammals and insects, odorants bind to receptors in the cilia or dendrites of olfactory receptor neurons (ORNs), each of which expresses only one type of receptor. ORNs that express the same receptors then converge onto the same glomerulus in olfactory bulb (mammals) or antennal lobe (insects), whose activity patterns contain the information about external odor stimuli [13, 17–19]. A key question that we want to address here is how ORNs best represent external olfactory information that can be interpreted by the brain to guide an animal’s behavior [4, 13, 20].

It has long been hypothesized that the input-output response functions of sensory neurons are “selected” by statistics of the stimuli in the organism’s natural environment to transmit a maximum amount of information about its environment, generally known as the efficient coding hypothesis [21, 22] or the related InfoMax principle [23, 24]. For instance, the contrast-response function of interneurons in the fly’s compound eye can be well approximated by the cumulative probability distribution of contrast in the natural environment [22]. The receptive fields of neurons in early visual pathway are thought to exploit statistics of natural scenes [25–29]. Similar result has also been observed in the auditory system [30]. In all these cases, to achieve maximum information transmission, an “ideal” neuron should transform the input distribution into a uniform output distribution [22, 31] and a population of neurons should decorrelate their responses [28, 32, 33].

However, unlike light or sound, which can be characterized by a single quantity such as wavelength or frequency, there are a huge number of odorants each with its own unique molecular structure and different physiochemical properties [34, 35]. The high dimensionality of the odor space thus poses a severe challenge for the olfactory system to code olfactory signals. Fortunately, typical olfactory stimuli are sparse with only a few types of odorant molecules in an odor mixture [1, 2, 36]. The sparsity of the odor mixture immediately reminds us of the powerful compressed sensing (CS) theory developed in computer sceince and signal processing community [37, 38]. The CS theory shows that sparse high dimensional signals can be encoded by a small number of sensors (measurements) through random projections; and the highly compressed signal can be reconstructed (decoded) with high fidelity by using an *L*_1_-minimization algorithm [37–39]. However, conventional CS theory assumes the sensors to have a linear response function with essentially an infinite dynamic range [40]. In contrast, ORN response is highly nonlinear [41, 42], with a typical dynamic range less than 2 orders of magnitude, which is far less than the typical concentration range of odorants [42].

The use of the CS theory has recently been explored in olfactory systems. For example, Zhang and Sharpee proposed a fast reconstruction algorithm in a simplified setup with binary ORNs and binary odor mixtures without concentration information [43]. In another work, Krishnamurphy *et al*. studied how the overall “hour-glass” (compression followed by decompression) structure of the olfactory circuit can facilitate olfactory association and learning, again with the assumption that ORN responses to odor mixtures are linear [44]. Following ideas in CS theory, Singh *et al* recently proposed a fast olfactory decoding algorithm that might be implemented in the downstream olfactory system [45]. However, All these studies primarily focus on the downstream decoding and learning of the compressed signals by assuming a linear neuron response function. The question of how ORNs with a nonlinear input-output response function can best compress the sparse high dimensional odor information remains unanswered.

Another related study is the recent work by Zwicker *et al*. [46] where the authors investigated the maximum entropy coding scheme for the olfactory system by using a simplified binary response function, where an odor only induces a response when its concentration is above a threshold that is inversely proportional to the receptor sensitivity to the odor. They found two conditions for the binary ORNs to maximize the information transmission. The first condition is that each ORN on average responses to half of the odors, i.e., half of the odors have a concentration that is higher than the corresponding threshold; and the other condition is that the responses from different ORNs need to be uncorrelated. These results were obtained by studying the average activities of the ORNs and their correlations. However, due to the limitation of the binary input-output response function and the specific prior for the sensitivity distribution used in [46], the optimal coding strategy for neurons with realistic physiological properties remains unclear.

To address these important open questions, here we study the optimal coding scheme by using a realistic ORN input-output response function where the ORN output depends on the odor concentration continuously in a nonlinear (sigmoidal with odor concentration on a logarithmic scale) form characterized by its sensitivity, or equivalently the inverse of the half-maximum response concentration. By optimizing the input-output mutual information in the full sensitivity matrix space without any prior and following general insights from compressed sensing for sparse odor mixtures, we systematically study the statistical properties of the optimal sensitivity matrix and their dependence on odor statistics.

We found that the optimal ORN sensitivity matrix is sparse, i.e., each receptor only responds to a fraction of the odorants in its environment and its sensitivity to the other odorants is *zero*. The sparsity itself is a robust (universal) feature of the optimal sensitivity matrix, and the value of the optimal sparsity depends on statistics of the odor mixture and the number of ORNs. By using a simple mean-field theory, we show that the sensitivity matrix sparsity is caused by the competition (trade-off) between enhancing multiple odor detection and avoiding odor-odor interference. Next, we demonstrate the advantages of the maximum entropy (information) coding scheme in two downstream “decoding” (learning) tasks: reconstruction and classification of odor stimuli. Finally, we generalize our theory to neurons with a finite basal activity, where we found that the optimal coding strategy is to allow co-existence of both odor-evoked inhibition and activation with the fraction of inhibitory interactions depending on the basal activity. Comparisons with existing data in different organisms are consistent with our theory, which provides a unified framework to understand olfactory coding. Possible predictions based on our theory and future directions to incorporate more biological complexities such as odor-odor and receptor-receptor correlations in our model are also discussed.

## II. RESULTS

We first describe the mathematical setup of the problem before presenting the results. An odor mixture can be represented as a vector ***c*** = (*c*_1_,…,*c_N_*), where *c_j_* is the concentration of odorant (ligand) *j* (= 1, 2,…,*N*) and *N* is the number of all possible odorants in the environment. A typical odor mixture is sparse with only *n*(≪ *N*) odorant molecules that have non-zero concentrations. As illustrated in Fig. 1(a) (dotted box), the odor mixture signal ***c*** is sensed by *M* sensors. The encoding process, which maps ***c*** to the ORN response vector ***r*** = (*r*_1_, *r*_2_, …,*r_M_*), is determined by the bipartite odorant-ORN interaction network characterized by the sensitivity matrix ***W***, whose elements are denoted as *W_ij_*, the sensitivity of *i*-th sensor (ORN) to *j*-th odorant for all odorant-ORN pairs with *j* = 1, 2,…,*N* and *i* = 1, 2,…, *M*. For simplicity, we used a simple competitive binding model [47, 48] [and see Supplemental Material (SM)], in which the normalized response of ORN *i* (=1, 2, ⋯, *M*) to odor ***c*** can be described by a nonlinear function

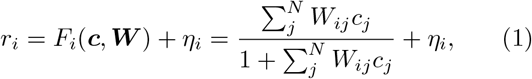

where *F_i_* is the response function which is taken to be the form in Eq. 1, and *η_i_* represents the noise term. For convenience, we assume *η_i_* is a Gaussian noise with zero mean and standard deviation *σ*_0_. Other forms of the nonlinear response function and noise can be used without affecting the general conclusions of our paper.

**FIG. 1.**
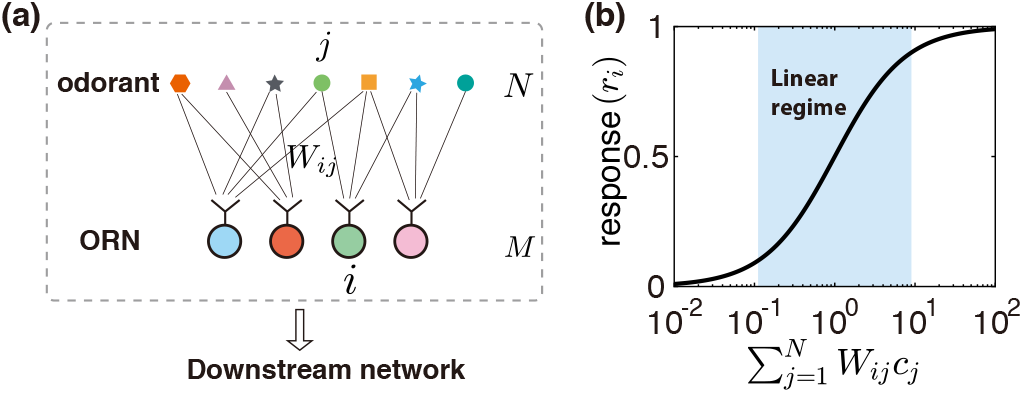
Schematics of peripheral odor coding. (a) Illustration of interaction between odorants and odor receptors. There are *N* possible odorant molecules and *M* ORNs. The interaction between the *i*’th ORN and the *j*’th odorant is characterized by the sensitivity *W_ij_*. The odorant-ORN interactions in the peripheral olfactory system (the dotted box) are characterized by the (*M* × *N*) sensitivity matrix ***W***. The odor information collected by the ORNs are passed on to downstream network for further processing. (b) Typical ORN-odorant dose-response curve according to Eq. (1). The range of linear response (in the log scale) is highlighted by the shaded area.

As illustrated in Fig. 1(b), the input-output response curve is highly nonlinear (Sigmoidal) resulting in a finite response range for each sensor, which is less than the range of concentration for a typical odorant molecule. Therefore, to encode the full concentration range of an odorant molecule, an odorant needs to interact with multiple sensors with different sensitivities. On the other hand, given the fact that *M* < *N*, each sensor has to sense multiple odorant molecules. These two considerations lead to the many-to-many odor-receptor interaction network characterized by the sensitivity matrix ***W*** = {*W_ij_* | *i* = 1, 2,…, *M; j* = 1, 2,…,*N*}.

Eq. (1) maps the external odor stimulus ***c*** to the internal neuronal activity ***r*** = (*r*_1_, *r*_2_, ⋯, *r_M_*). The downstream olfactory circuits then use this response pattern to evaluate (decode) odor information (both their identities and concentrations) in order to guide the animal’s behaviors. The quality of encoding odor information by the periphery ORNs directly sets the upper limit of how well the brain can decode the odor information [49]. In this paper, we focus on discovering the key statistical properties of the sensitivity matrix ***W*** that allows biologically realistic ORNs to best represent the external odor information.

A given odor environment can be generally described by a probability distribution *P*_env_(***c***). To convey maximum information about external odor stimuli in their response patterns to the brain, ORs/ORNs can adjust their sensitivity matrix ***W*** to “match” odor statistics *P*_env_(***c***). Without any assumption on what information the brain may need, the mutual information *I*(***c, r***) between stimuli and response pattern of ORNs sets the limit on how much odor information are received by the peripheral ORNs and thus serves as a good “target function” to be maximized [23, 24, 46, 50–52]. *I* is defined as

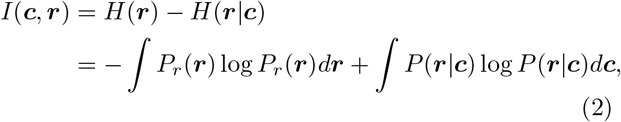

where *H*(***r***) and *H*(***r***|***c***) are the entropy of output distribution *P_r_*(***r***) and conditional distribution *P*(***r***|***c***). In the limit of small noise, the second term is independent of ***W*** and negligible, hence, we will use *H*(***r***) as our target function for optimization.

*H*(***r***) depends on ***W*** and *P*_env_(***c***) because *P*_r_(***r***) depends on ***W*** and *P*_env_(***c***):

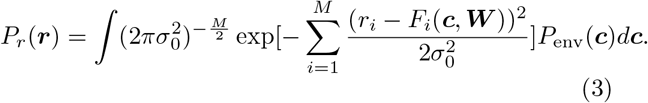

In this paper, we study the optimal coding strategy by maximizing the mutual information *I* or equivalently the differential entropy *H* with respect to the sensitivity matrix ***W*** for different odor mixture statistics *P*_env_(***c***) and different numbers of ORNs. The mutual information as given in Eq. (2) can only be computed analytically for very simple cases. For more general cases, we used the covariance matrix adaptation - evolution strategy (CMA-ES) algorithm to find the optimal sensitivity matrix [53, 54] (see SM for technical details).

### A. The optimal sensitivity matrix is sparse

Odor concentration varies widely in natural environment [1, 55]. To capture this property, we studied the case where the odorant concentrations in an odor mixture follows a log-normal distribution with variance 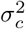. Other broad distributions such as power law distributions are also studied without changing the general conclusions. For simplicity, we consider the case where odorants appear independently in the mixture; more realistic consideration such as correlation among odorants will be discussed later in the Discussion section.

For given odor statistics (characterized by *N, n* and *σ_c_*), and a given number of nonlinear sensors *M*, we can compute and optimize the input-output mutual information *I*(***W***|*N, n, σ_c_; M*) with respect to all the *M* × *N* elements in the sensitivity matrix ***W***. We found that the optimal sensitivity matrix ***W*** is “sparse”: only a fraction (*ρ_w_*) of its elements have non-zero values [sensitive, shown as the colored elements in Fig. 2(a)], and the rest are insensitive [the black elements in Fig. 2(a)], with essentially zero values of *W_ij_*. The sparsity in the optimal ***W*** was not found in the previous theoretical study [46] mainly due to the oversimplified binary ORN response function used there. From the histogram of ln(*W_ij_*) shown in Fig. 2(b), it is clear that elements in the optimal sensitivity matrix fall into two distinctive populations: the insensitive population that has practically zero sensitivity [note the log scale used in Fig. 2(b)], and a sensitive population with a finite sensitivity. For the cases when the odor concentration follows a log-normal distribution, the distribution of the sensitive (nonzero) elements *P_s_*(*w*) can be fitted well with a log-normal distribution as shown in Fig. 2(b).

**FIG. 2.**
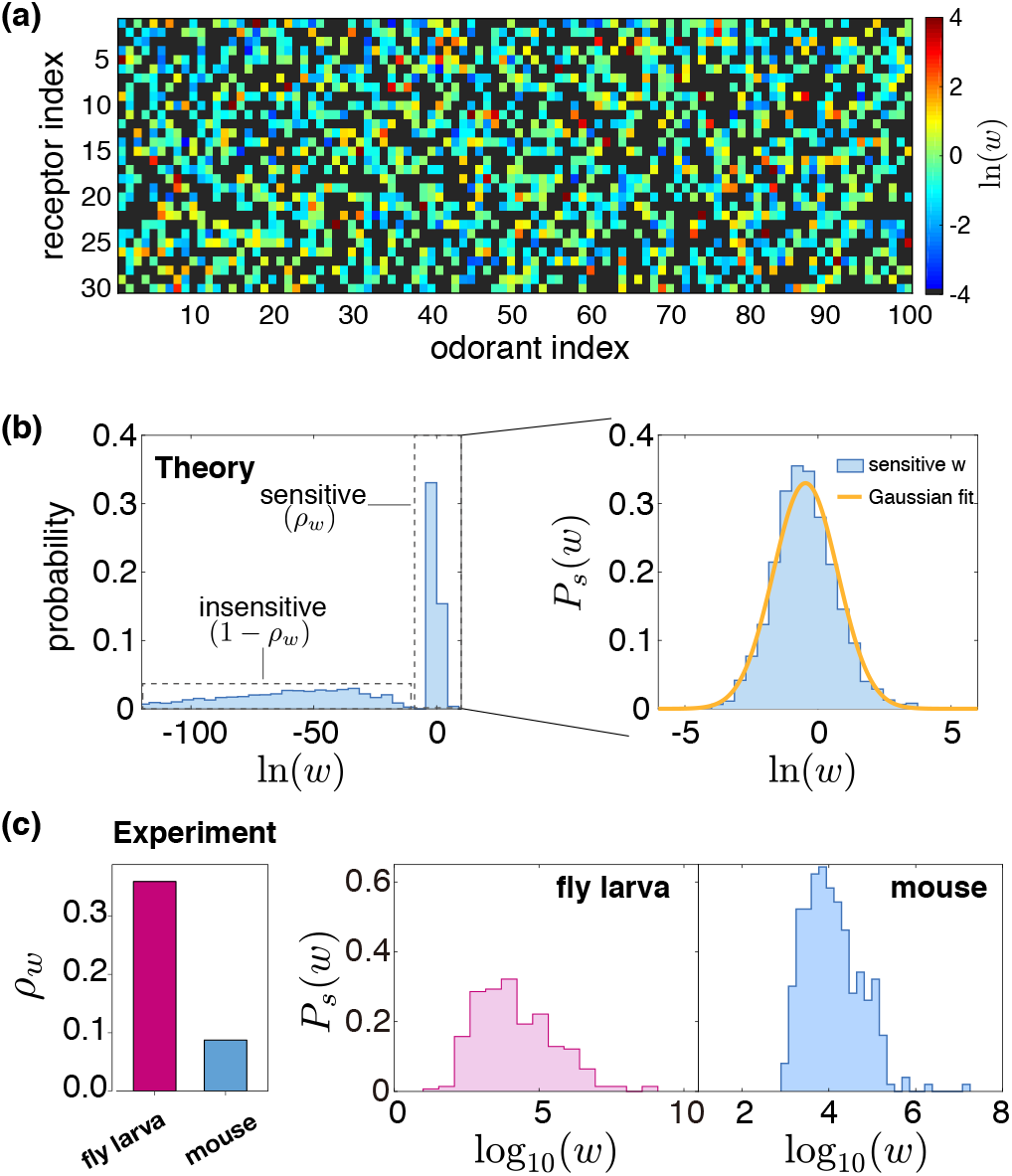
Statistics of the optimal sensitivity matrix elements from theory and comparison with experiments. (a) Heatmap of a typical optimal sensitivity matrix from our model. Color indicates the value of ln(*W_ij_*), black indicates the “inactive” or negligible interactions. (b) Histogram of all the *W_ij_* values from our model. It shows a bimodal distribution: an insensitive part with near zero *W_ij_* and a sensitive part with nonzero *W_ij_*. The distribution of the sensitive elements can be fitted by a log-normal distribution. (c) Experimental data from fly larva and mouse. Left: the fraction of sensitive odorant-receptor interactions *ρ_w_* estimated in experiments for fly larva [42] and mouse [56]. Right: the histogram of sensitive *W_ij_, P_s_*(*w*), for fly larva and mouse. Model parameters are *N* = 100, *M* = 30, *n* = 2, *σ_c_* = 2, *μ* = 0.

Our main finding here, i.e., sparsity in the odor-receptor sensitivity matrix, is supported by existing experimental measurements. As shown in Fig. 2(c), the sparsity *ρ_w_* is estimated to be ~ 0.4 for fly larva [42] and ~ 0.1 for mouse [56]. Additionally, the broad distribution of the non-zero sensitivity observed in our model also agree qualitatively with those estimated from experiments [Fig. 2(c), two right panels], which are slightly skewed log-normal distributions.

Besides the distribution of the individual sensitivity matrix elements, we also calculated the row (sensor)-wise and column (odorant)-wise rank-order correlation coefficients (Kendall’s tau, *τ*) and compared them with those from the same matrix but with its elements shuffled randomly. As shown in the Supplemental Material (SM), we found that both the rows and columns (Fig. S1 in SM) in the optimal matrix have a higher level of orthogonality (and thus independence) than that from random matrices. This orthogonality in the optimal ***W*** matrix leads to a higher input-output mutual information than those from the shuffled matrices [see Fig. S2(a) in SM] and a nearly uniform distribution of ORN activity for different odor mixtures [see Fig. S2(b)-(d) in SM].

### B. The optimal sparsity depends on odor statistics and the number of sensors

The statistics of the optimal sensitivity matrix elements are characterized by the sparsity *ρ_w_* defined as the fraction of *non-zero* elements in ***W***, and the distribution of the sensitive (nonzero) elements, *P_s_*(*w*), which is further characterized by its mean (*μ_w_*) and standard deviation (*σ_w_*). Note that the sparsity parameter *ρ_w_* is defined in such a way that a smaller value of *ρ_w_* corresponds to a sparser sensitivity matrix. We investigated systematically how *ρ_w_, μ_w_*, and *σ_w_* depend on statistical properties of the odor mixture characterized by *N, n*, and *σ_c_*, as well as *M*, the total number of sensors (ORNs).

We found that as the odor concentration becomes broader with increasing *σ_c_, ρ_w_* increases [Fig. 3(a)]. This is expected as more receptors with different sensitivities are required to sense a broad range of input concentrations. When we increased the odor mixture sparsity *n* or the total number of possible odors *N*, the optimal sensitivity matrix sparsity *ρ_w_* decreases [Fig. 3(b)&(c)]. In general, as the mapping from odor space to ORN space becomes more “compressed” with larger values of *n* and/or *N*, the optimal strategy is to have each receptor respond to a smaller fraction of odorants to avoid saturation. Finally, we gradually increased the number of receptors *M* with fixed values of *N, n*, and *σ_c_*. We found that *ρ_w_* decreases, i.e., the sensitivity matrix becomes more sparse as the number of sensors *M* increases [Fig. 3(d)]. This somewhat counter-intuitive result can be understood as the system has more sensors to encode signals, each sensor can respond to a smaller number of odors to avoid interference. For all the cases we studied, when the odor concentrations follow a log-normal distribution, then the distribution of the non-zero sensitivities in the optimal sensitivity matrix follows roughly a log-normal distribution with its mean *μ_w_* and standard deviation *σ_w_* depending on the odor statistics (*σ_c_, n, N*) and the number of ORNs *M* (see Fig. S3 in SM).

**FIG. 3.**
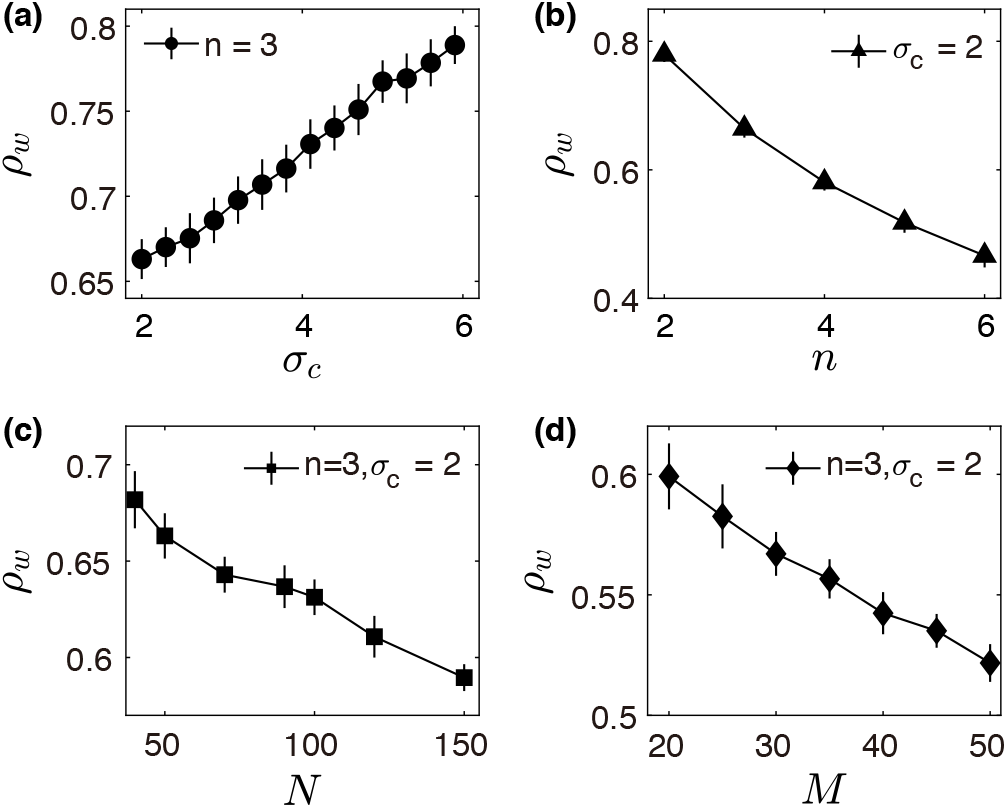
Dependence of *ρ_w_* on the width of odor log concentration *σ_c_* (a), the input sparsity *n* (b), the number of total odorants *N* (d), and the number of receptors *M* (d). In (a-b), *N* = 50, *M* = 13, in (c), *M* = 13, and in (d), *N* = 50. Error bars are standard deviation of 40 times simulation.

To verify whether sparsity is a general (robust) feature in the optimal sensitivity matrix, we studied the cases when the odor concentration follows different distributions, such as a symmetrized power-law distribution, *P*_env_(*c*) ∝ exp(−*β*| ln *c*|)[see Fig. S4(a) in the SM for the comparison with log-normal distribution], with different exponent *β*. For all values of *β* studied, there is always a finite sparsity *ρ_w_* < 1 in the optimal sensitivity matrix. As shown in Fig. 4(a), *ρ_w_* decreases slightly when *β* increases and the odor concentration distribution becomes narrower, which is consistent with the previous cases when the odor concentration distribution is log-normal [Fig. 3(a)]. However, as shown in Fig. 4(b), the distribution of the sensitive elements, *P_s_*(*w*), does not follow exactly a log-normal distribution [see Fig. S4 (b) in SM]. In fact, *P_s_*(*w*) is asymmetric in the ln(*w*) space with a skewness that depends on *β* as shown in the inset of Fig. 4(b).

**FIG. 4.**
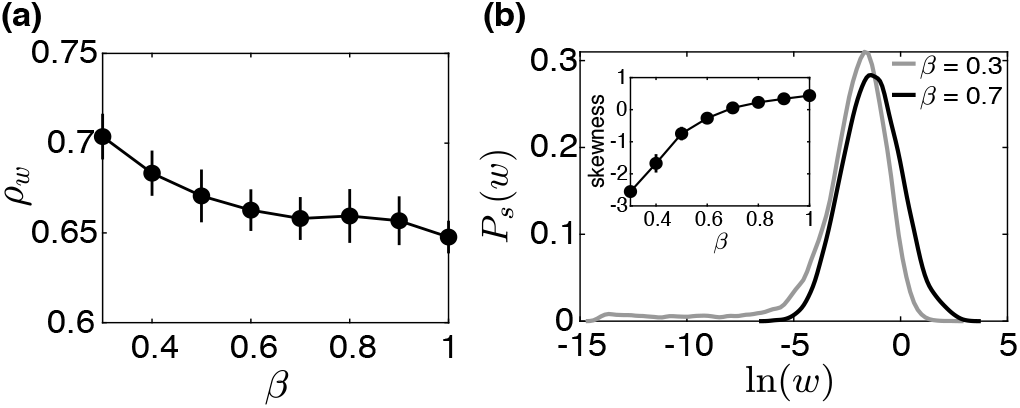
The optimal sensitivity matrix for the symmetric power-law odor concentration distribution *P*_env_(*c*) ∝ exp[−*β*| ln *c*|]. (a) The sparsity *ρ_w_* versus the power-law exponent *β*. (b) The distribution of the non-zero sensitivities *P_s_*(*w*) for *β* = 0.3, 0.7. Inset shows the dependence of the skewness of the distribution on *β*. Parameters: *N* = 50, *M* = 13, *n* = 3, *σ*_0_ == 10^−3^.

Taken together, our results suggest that sparsity in the sensitivity matrix is a robust feature for nonlinear compressed sensing problems. This theoretical finding is supported by and explains existing experiments in olfactory systems [42, 56]. Our study also showed that the nonzero sensitivities follow a broad distribution whose exact shape, mean, and variance depend on odor statistics and total number of ORNs.

### C. The origin of sparsity in the optimal sensitivity matrix

Given the constraint that the number of sensors is much smaller than the possible number of odorants, i.e., *M* ≪ *N*, each sensor needs to respond (sense) to multiple types of odorant molecule so that all odorant molecules can be sensed by at least one sensor. However, in an odor mixture with a few types of odorant molecules, two or more odorants in the mixture can bind with the same sensor and interfere with each other, e.g., by saturating the nonlinear sensor. The probability of interference increases with the sparsity of the sensitivity matrix. This tradeoff between sensing multiple odorants and the possible interference determines the sparsity in the optimal sensitivity matrix. We demonstrate this tradeoff and its effect more rigorously by developing a mean-field theory (MFT) as described below.

We begin with the simplest case where many receptors sense only one odorant (*N* = 1, *M* ≫ 1) where obviously there is no interference. As first proposed by Laughlin [22], the optimal coding scheme is for the *M* receptors to distribute their sensitivities according to the input concentration distribution so that the output distribution is uniform. For the case when the distribution of the odorant concentration is log-normal with a standard deviation *σ_c_*, the optimal sensitivity distribution *P*_1_(*w*) that maximizes *H*(***r***) is also approximately a log-normal distribution:

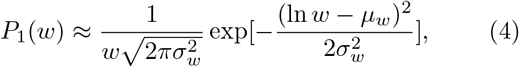

where the mean *μ_w_* = 0 and the variance 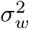 increases with the variance 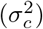 of logarithmic concentration distribution. More importantly, we show analytically that in general the coding capacity *I*_1_ increases logarithmically with the number of receptors M when *M* ≫ 1 (see SM for details), which is verified by simulation results as shown in Fig. 5(a):

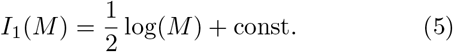

**FIG. 5.**
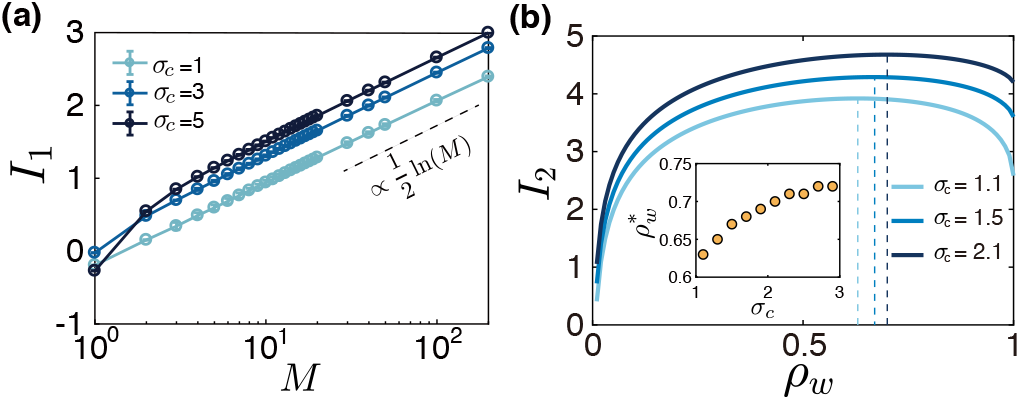
The tradeoff between increasing single odorant information and interference among multiple odorants. (a) The differential entropy with one odorant, *I*_1_, versus the number of receptors *M* for different width (*σ_c_*) of odor log-concentration distribution. *I*_1_ increases monotonically with *M* but it only grows logarithmically with *M* for large *M* (dashed line). (b) Differential entropy *I*_2_ for the case with two odorants in the mixture with their concentrations following the same log-normal distribution with width *σ_c_. I*_2_ depends non-monotonically on the fraction of sensitive receptors *ρ_w_*(= *m*/*M*) with a maximum (marked by the dashed lines) at 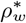 that depends on *σ_c_*, which is shown in the inset.

This means that sparsity *ρ_w_* = 1, i.e., all sensitivities should be nonzero because there is no interference when only one type of odorant molecule (*n* = 1) is present in the mixture. However, it is important to note that the maximum mutual information only increases weakly (logarithmically) for large *M*.

We next consider the case where two odorants are sensed by multiple receptors (*N* = 2, *M* ≫ 1). Let’s denote the number of receptors that respond to each odorant as *m*(*m* ≤ *M*) and the sparsity *ρ_w_* = *m*/*M*. If each odorant is sensed by a disjoint set of receptors, the total differential entropy will simply double the amount for a single odorant: *I*_2_(*m*) = 2*I*_1_(*m*). However there is a finite probability *p* = *m*/*M* = *ρ_w_*, that a given receptor in one set will also respond to the other odorant. Therefore, on average there are *m* × *p* = *m*^2^/*M* receptors whose output is “corrupted” due to interference between two different odorants in a given mixture. We can write down the differential entropy as

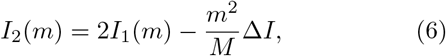

where *I*_1_(*m*) is the maximum differential entropy for one odor [Eq. (5)] and Δ*I* is the marginal loss of information (entropy loss), which can be approximated by Δ*I* ≈ *α*(*I*_1_(*m* + 1) − *I*_1_(*m*)) ≈ *α∂I*_1_(*m*)/*∂m* where *α* ≤ 1 is the average fraction of information loss for a “corrupted” sensor. We can then obtain the optimal value of *m* by maximizing *I*_2_(*m*) with respect to m. For *m* ≪ *M*, the interference effect is small, so *I*_2_(*m*) ≈ 2*I*_1_(*m*), which increases with *m* logarithmically according to Eq. (5). As *m* increases, the interference effect given by the second term on the RHS of Eq. (6) increases with *m*, which is faster than the slow logarithmic growth of 2*I*_1_(*m*). This leads to a peak of *I*_2_(*m*) at an optimal value of *m* = *m*\* < *M* or a sparsity of the sensitivity matrix *ρ_w_* = *m*\*/*M* < 1 [Fig. 5(b)].

In the MFT, we can compute the olfactory coding and interference by ignoring the weak rank-order correlation in the optimal sensitivity matrix and assuming the distributions for the optimal sensitivity matrix elements are *i.i.d*. In particular, we used the following approximation for the distribution of the sensitivity matrix ***W***:

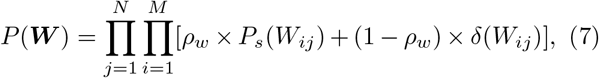

where *ρ_w_* is the matrix sparsity, and *P_s_*(*W_ij_*) is a smooth distribution function, which is approximated here as a log-normal distribution with mean *μ_w_* and standard deviation *σ_w_* as given in Eq. (4). The mean differential entropy of ORN response pattern

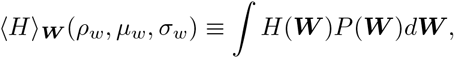

which is averaged over the distribution of the sensitivity matrix ***W***, can be maximized with respect to the parameters *ρ_w_, μ_w_*, and *σ_w_* (see SM for details). The resulting optimal parameters agree with our direct numerical simulations qualitatively with a sparsity *ρ_w_* < 1 that increases with the width of the input distribution *σ_c_* (see Fig. S5 in SM).

### D. The optimal sparse sensitivity matrix enhances downstream decoding performance

The response patterns of ORNs form the internal representation of external odor stimuli that the higher (downstream) regions of the brain can use to infer the odor information for controlling the organism’s behavior. In previous sections, we focused on understanding the statistical properties of the optimal sensitivity matrix *W* that maximize mutual information between odor input and ORN output. Here in this section, we test whether the optimal sensitivity matrix can enhance the downstream decoding performance by examining two specific learning tasks: classification and reconstruction.

#### Task I: classification

The goal of the classification task is to infer the category of odor mixture such as the odor valence by training with similar odor stimuli. Classification is believed to be carried out by the *Drosophila* olfactory circuit, which we describe briefly here. After odor signals are sensed by ~ 50 ORNs, they are relayed by the projection neurons (PNs) in antennal lobes to a much larger number of Kenyon cells (KCs) in the mushroom body (MB), as illustrated in Fig. 6(a). Each of the ~ 2000 KCs in MB on average receives input from ~ 7 randomly selected PNs [57]. A single GABAergic neuron (APL) at each side of the brain can be activated by the KCs and inhibits KCs globally [58]. Such random expansion and global inhibition enable sparse and decorrelated representation of odor information in MB [57, 59, 60]. The large number of KCs in MB then converge to a few (only 34) mushroom body output neurons (MBONs) [61], which project to the other brain regions and drives attractive or repulsive behavior [62]. Olfactory learning is mainly mediated by the dopaminergic neurons (DANs) which controls the synaptic weights between KCs and MBONs [63].

**FIG. 6.**
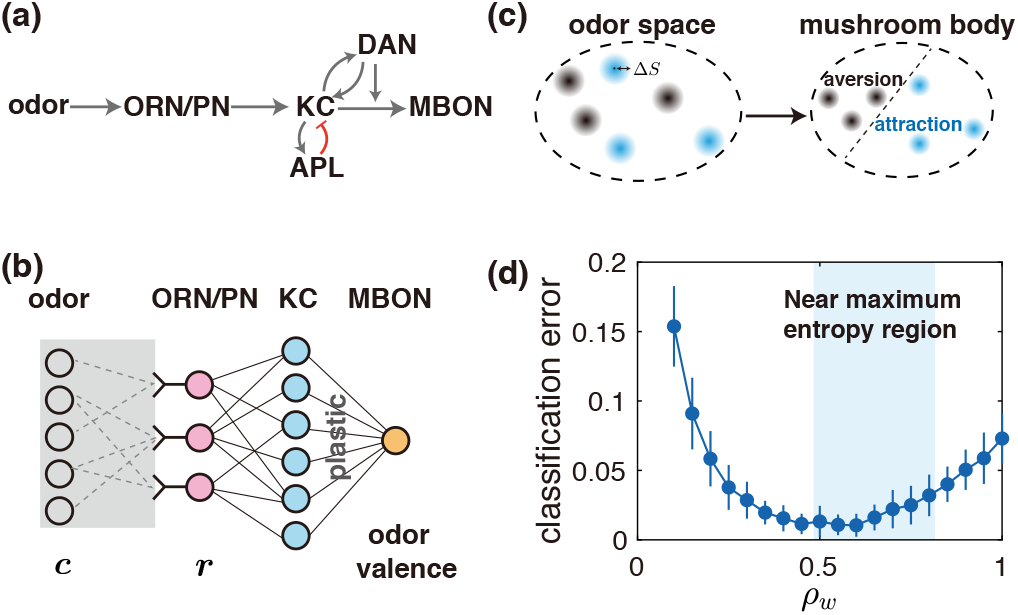
Maximum entropy coding facilitates olfactory learning and classification. (a) Schematics of the neural circuitry for olfactory learning in fly, see text for detailed description. (b) A simplified model of the fly olfactory system shown in (a) for learning the valence of odor stimuli, where the effect of dopaminergic neurons (DAN) is replaced by simple plastic weights from KC to MBON. (c) Odors are organized as clusters of size Δ*S* and are randomly assigned with an odor valence. The decoding network receives the response pattern of ORNs and classifies them into the right categories. 100 clusters were drawn and each cluster contains 50 variations, resulting in 5000 odor stimuli among which 80% were used as training data and the rest were used as testing data. (d) Classification performance with respect to the sparsity of sensitivity matrix. Best performance appears at around *ρ_w_* = 0.6, within the 95% maximum entropy region. Parameters: *N* = 100, *M* = 10, *n* = 3, *σ_c_* = 2, *σ*_0_ = 0.05, Δ*S* = 0.1, 500 KC units and two odor categories. Error bars are standard deviation from 40 simulations.

To mimic the properties of MB, our model “classifier” network, as illustrated in in Fig. 6(b), contains a high dimensional mixed-layer (KCs). For simplicity, we consider a single readout neuron. Each KC unit in the mixed-layer pools the ORNs with a fixed random, sparse matrix. Only the synaptic weights from the KCs to the readout neuron are plastic. To consider the variability of natural odors, we assumed that odor stimuli fall into clusters whose centers represent corresponding typical odor stimuli. Members in a given cluster are variations to the centroid [64]. The radius of a cluster Δ*S* characterizes the variability of a specific odor mixture. Centroids were drawn from *P*_env_(***c***) with randomly assigned labels (attractive or aversive). Members inside each cluster were generated by adding noise of size Δ*S*, which results in clouds of points in the odor space [Fig. 6(c)] with each cloud having a randomly assigned label (see SM for details).

The synaptic weights from the KCs to the readout neuron are trained by using a simple linear discriminant analysis (LDA) method although other linear classification algorithms such as support vector machine (SVM) would also work. After training, the performance of the “classifier” is quantified by the accuracy of classification on the testing dataset.

To test effects of different coding schemes on the classification performance, we vary the distribution of the sensitivity matrix elements by changing the sparsity *ρ_w_* without changing the distribution of the non-zero sensitivity matrix elements (e.g., the log-normal distribution with fixed mean and variance). The output of the coding process ***r***(***c, W***) serves as the input for the “classifier” network and the classifier error is computed for different values of *ρ_w_*. As shown in Fig. 6(d), we find that the best performance is achieved near *ρ_w_* = 0.6, which belongs to the range of *ρ_w_* with large mutual information between odor input and the ORN/PN response [shaded region in Fig. 6(d)]. Changing parameters such as *M, n* and number of categories give similar results (see Fig. S6 in SM). In line with recent studies which show that sparse highdimensional representation facilitates downstream classification [64, 65], our results suggest that maximum entropy coding at the ORNs/PNs level may enhance classification by retaining maximum odor mixture information in a form that can be decoded by the KCs through random expansion.

#### Task II: reconstruction

This goal of the reconstruction task is to infer (decode) both the composition and the exact concentrations of all odorant components in an odor mixture from the sensor responses. This more stringent task is motivated directly by the original compressed sensing problem in computer science, its relevance to the olfactory systems will be discussed later in the Discussion section.

As illustrated in Fig. 7(a), the output of the coding process ***r***(***c, W***) serves as the input for the downstream reconstruction network. Here, we used a generic feedforward artificial neural network (ANN) with a few (1-5) hidden layers and a output layer that has the same dimension *N* as the odor space. We trained the ANN with a training set of sparse odor mixtures drawn from the odor distribution *P*_env_(***c***), and tested its performance by using new odor mixtures randomly drawn from the same odor distribution. Denote the reconstructed odor vector as ***ĉ*** and a binary vector ***ξ*** associated with ***c***, i.e., *ξ_i_* = 1 if *c_i_* ≠ 0, otherwise, *ξ_i_* = 0. Due to the sparse nature of odor mixture and the wide concentration range, the reconstruction error 𝓛 is defined as the sum of “identity error” 𝓛_1_ and “intensity error” 𝓛_2_:

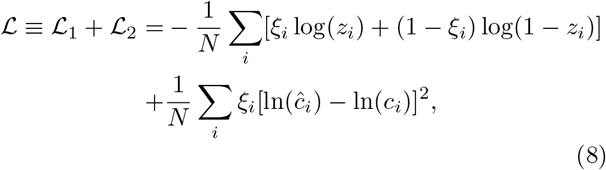

where 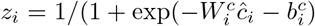 with 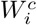 and 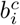 determined from training (supervised learning).

**FIG. 7.**
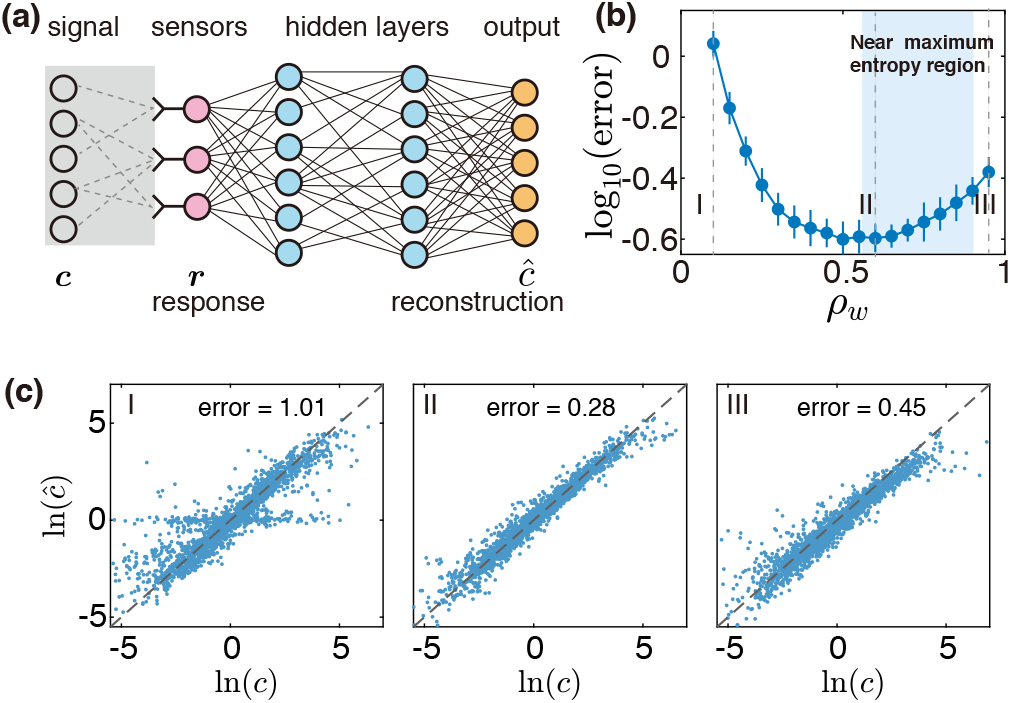
Maximum entropy coding facilitates signal reconstruction. (a) Schematic of the reconstruction task. The decoding feed-forward network with multiple hidden layers (blue nodes) receives the response ***r*** of the peripheral sensors (red nodes) as ‘input’ and produces the output ***ĉ*** to reconstruct (infer) the original signal ***c***. (b) The minimum reconstruction error on the testing set is achieved around sparsity level *ρ_w_* = 0.6 of ***W***, within the 95% maximum entropy region (shaded area).(c) Comparison of the original *c_i_* and reconstructed *ĉ_i_* for three different sparsity of ***W***: I. *ρ_w_* = 0.1, II. *ρ_w_* = 0.6, III. *ρ_w_* = 0.95, which are labeled by the dashed lines in panel (b). Parameters used are: *N* = 100,*M* = 20,*n* = 2,*σ_c_* = 2, *σ*_0_ = 0.05, two hidden layers, each with 100 units. Error bars in (b) are standard deviation from 40 times simulation.

The reconstruction error depends on the coding matrix ***W***, in particular its sparsity *ρ_w_*, as shown in Fig. 7(b). Pair-wise comparisons of non-zero concentrations in the original and reconstructed odor mixtures for three different coding regimes are shown in Fig. 7(c) (see Fig. S7 in SM for a direct comparison of the whole reconstructed and original odor vectors).The best performance is achieved around *ρ_w_* = 0.6, within the region where sparse ***W*** enable nearly maximum entropy coding (shaded region), this property is insensitive to the number of hidden layers in the reconstruction network (see Fig. S8 in SM). Our results show that the optimal entropy code provides an efficient representation of the high-dimensional sparse data so that the downstream machine learning algorithms can achieve high reconstruction accuracy.

### E. Optimal coding strategy for ORNs with a finite basal activity

So far, we have only considered the case where the neuron activity is zero in the absence of stimulus and odorants only activate the ORs/ORNs. It has been widely observed that some ORNs show substantial spontaneous activities, and some odorants can act as inhibitors to suppress the activities of neurons they bind to [41, 66, 67], as shown in Fig. 8(a). The presence of an inhibitory odorant can shift a receptor’s dose-response curve to an excitatory odorant, thereby diminishing the sensitivity of the receptor to excitatory odorants[68]. It is then natural to ask what is the optimal design of the sensitivity matrix to maximize coding capacity if odorants can be either excitatory or inhibitory. To answer this question, we used a two-state model to characterize both odor-evoked excitation and inhibition [68]. Now, the interaction between the odorant *j* and ORN *i* has two possibilities – it can be either excitatory with a sensitivity 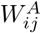 or inhibitory with a sensitivity 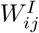. The normalized response of *i*-th ORN to odor mixture ***c*** is

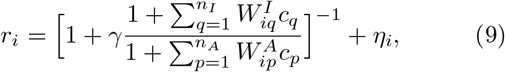

where *γ* determines the basal activity by *r*_0_ = 1/(1 + *γ*), *n_A_* and *n_I_* are the number of excitatory and inhibitory odorants to the *i*-th receptor, and *η_i_* is a small Gaussian white noise.

**FIG. 8.**
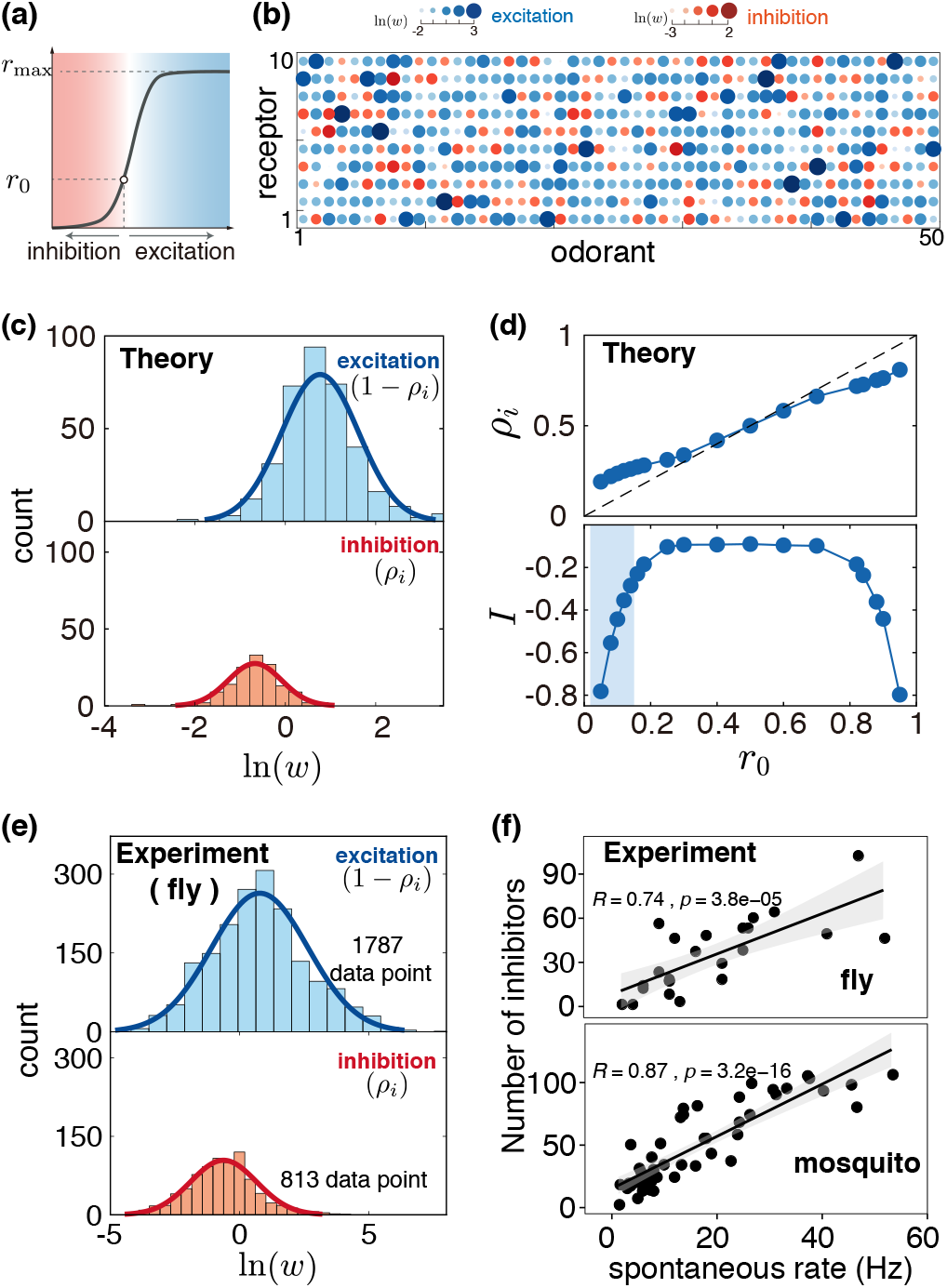
The optimal sensitivity matrix for ORNs with a finite basal activity and comparison with experiments. (a) Schematic of ORN response to excitatory (blue region) and inhibitory odorants (red region). Note that the neuron has a finite acitivity *r*_0_ in the absence of any stimulus. (b) Heatmap of a typical optimal *W* from our model, with the size of the elements indicating the strength of excitatory (blue) and inhibitory (red) interactions. (c) Both the excitatory and inhibitory interactions in optimal *W* can be well approximated by log-normal distributions (solids lines). (d) The fraction of inhibitory interaction *ρ_i_* increases the basal activity *r*_0_ nearly linearly (upper panel). The differential entropy *I* also increases with *r*0 (lower panel). The shaded region shows the range of *r*_0_ corresponding to the fraction of inhibitory interaction estimated from experiments [41], which coincide with the range of *r*_0_ where the differential entropy increases sharply with *r*_0_. (e) The distributions of the estimated relative excitatory and inhibitory receptor-odor sensitivities from experimental data for the fly [41]. Both distributions can be well fitted by log-normal distributions. (f) The correlation between the number of odorants that inhibit an ORN with the ORN’s spontaneous activity obtained from experimental data on fly [41] (upper panel) and mosquito [69] (lower panel). Each point corresponds to an ORN, the line is the linear fit and shaded region is the 0.95 confidence interval. Model parameters used are *N* = 50, *M* = 10, *n* = 2, *σ_c_* = 2, with 40 repeated simulations. In (d), error bars are small, comparable to the size of the symbols. In (b) and (c), *r*_0_ = 0.18.

Our simulations show that with a finite spontaneous activity, the receptor array achieves maximum entropy coding by assigning a certain number of inhibitory interactions in the sensitivity matrix [Fig. 8(b)]. The strength (sensitivity) of both the excitatory and inhibitory elements follow (approximately) log-normal distributions [Fig. 8(c)]. The fraction of inhibitory interaction (*ρ_i_*) in the optimal *W* is roughly proportional to the spontaneous activity of ORN *r*_0_, with only a slight deviation when *r*_0_ → 0 and *r*_0_ → 1 [Fig. 8(d, upper panel)]. Interestingly, as *r*_0_ → 0, *ρ_i_* approaches a finite value that is related to the fraction of zero sensitivity elements (1 − *ρ_w_*) we studied in the previous sections for ORNs without a spontaneous activity. As the basal activity increases, the coding capacity increases rapidly at first and quickly plateaus around *r*_0_ = 0.3 [Fig. 8(d, lower panel)]. The increase of coding capacity can be understood intuitively by considering that the effective dynamic range of receptors increases in the presence of inhibition. Odor-evoked inhibition enables receptors to work bi-directionally and avoid saturation when responding to many odorants simultaneously.

To verify our theoretical results, we have analyzed the statistics of the sensitivities for the excitatory and inhibitory interactions obtained from the experimental data in fly by Hallem and Carlson [41] as well as in mosquito by Carey *et al* [69]. As shown in Fig. 8(e) for the fly data, both the excitatory and inhibitory sensitivities follow log-normal distributions, which are consistent with our model results shown in Fig. 8(c). The mosquito data shows very similar results (see Fig. S9 in SM). Our theory also showed that the fraction of inhibitory interaction *ρ_i_* increases with the basal activity *r*_0_ as shown in Fig. 8(d, upper panel). We have tested this theoretical result from the experimental data. As shown in Fig. 8(f), the number of inhibitory odor-ORN interaction for an ORN shows a strong positive correlation with its basal activity for both fly and mosquito, which is in agreement with our theoretical prediction. Finally, we note that the relative basal activity 〈*r*_0_〉 from the experimnetal data [41] is smaller than 0.16 (see SM for detailed analysis), where the differential entropy rises sharply with *r_0_* as highlighted by the shaded region in Fig. 8(d, lower panel). Although an even higher spontaneous activity towards *r*_0_ = 0.5 can further increase the coding capacity, the gain is diminishing, while the metabolic cost increases drastically in maintaining the spontaneous activity [70]. Thus, an optimal basal activity would be expected in the shaded region of Fig. 8(d) due to the tradeoff between coding capacity and energy cost.

## SUMMARY AND DISCUSSIONS

In this paper, we studied how a relatively small number of nonlinear sensors (ORNs) with a limited dynamic range can optimize the transmission of high dimensional but sparse information in the environment. We found that the optimal sensitivity matrix elements follow a bi-modal distribution. For neurons without a basal activity, the sensitivity matrix is sparse – a neuron only responds to a fraction *ρ_w_*(< 1) of odorants with its sensitivities (to these odorants) following a broad distribution and it is insensitive to the rest (1 − *ρ_w_*) fraction of the odorants. This sparsity in the odor-ORN sensitivity matrix is a direct consequence of the finite dynamic range of the realistic nonlinear ORNs, which are different from the linear sensors in the conventional compressed sensing problem. For neurons with a finite basal activity *r*_0_, the optimal sensitivity distribution is also bi-modal with a fraction *ρ_i_* of the odor-neuron interaction inhibitory and the rest (1 − *ρ_i_*) fraction of the odor-neuron interaction excitatory and *ρ_i_* increases with *r*_0_. Details of the odor-receptor sensitivity distribution depend on the odor mixture statistics and the sensor characteristics, but the bi-modal distribution is robust. By investing the effects of different coding schemes on the downstream decoding/learning tasks, we showed that the maximum entropy code (representation) of the external signal enhances the performance of downstream reconstruction and classification tasks.

### Connection to experiments and testable predictions

Our primary finding - the sparsity in the odor-receptor sensitivity matrix ***W*** - seems to be consistent with existing experimental measurements of receptor-odor sensitivity matrices in different organisms [Fig. 2(c)]. Although the natural odor environment varies for different organisms, it is interesting to see that the broad distribution of the non-zero sensitivity observed in our model is consistent with the sensitivity matrices estimated from experiments in fly larva, mouse, adult fly, and mosquito [Fig. 2(c) and Fig. 8(e)&(f)]. The optimal coding strategy, if exists, would be the result of evolution. Thus, our theory may be tested by comparing olfactory systems in different species. In particular, our theory predicts that the sparsity parameter *ρ_w_* decreases with the number of ORNs *M* [Fig. 3(d)], which can be tested by measuring the sparsity in the odor receptor sensitivity matrices in different organisms.

The relatively high level of spontaneous activity in ORNs has long been thought to only play a role in the formation of topographic map during development [71]. A recent study shows that odor-evoked inhibition can code olfactory information that drives the behavior of the fly [68]. Our results provide a quantitative explanation for the advantage of having certain level of spontaneous basal activity and odor-evoked inhibitions in odor coding. For neurons with a finite basal activity, our theory predicts that the fraction of odorants that inhibit the neuron increases with the basal activity of the neuron. The data from adult fly and mosquito are consistent with this prediction [Fig. 8(f)]. However, powerful high throughput techniques such as Calcium imaging, which only indirectly measure the odor-ORN interaction, seem to be incapable of detecting inhibitory interaction [42]. Therefore, more large scale direct measurements using electrophysiological methods such as those done for *Drosophila* [41, 72] and mosquito [69] should be carried out to test our predictions in different organisms.

By considering how the coding capacity of ORNs changes with basal activity [Fig. 8(d)] and the associated extra energy cost [70], one can hypothesize the existence of an “optimal” *r*_0_. Our result suggests that as the number of sensors increases, the benefit of having basal activity diminishes, hence, the “optimal” *r*_0_ should decrease as the number of sensory neurons increases. Indeed, this is consistent with the fact that *E. coli* has 5 chemoreceptors [73] which work bi-directionally with a high basal activity *r*_0_ ≈ 1/3 − 1/2 [74], and *r*_0_ in mouse is smaller than that in fly [67]. Of course, more experiments across different organisms with different numbers of sensory neurons are needed to test this hypothesis.

### Future Directions

In this study, we assumed that odor information is contained in the instantaneous spiking rate of ORNs, and did not consider adaptation dynamics. Although adaptation plays an important role in all sensory systems [75], it happens in a relatively slower time scale than the time required for animals to detect and respond to odor stimuli [76, 77]. In general, sensory adaptation shifts the response function of the sensory neuron according to the background stimulus concentration and it leads to a larger but still finite effective dynamic range without changing the qualitative characteristics of the input-output response curve [75, 78]. Therefore, even though ORN level adaptation can further increase coding capacity at a slightly longer time scale as shown recently by Kadakia and Emonet [79], we do not expect it to qualitatively affect the optimal coding strategy found here. It remains an interesting question to understand how neuronal dynamics such as adaptation can be used for coding time-dependent odor signals.

We have used reconstruction and classification as two learning tasks to demonstrate the advantage of having maximum entropy coding at the ORN level. While the classification task has clear biological relevance, it is unclear to what extent animals need to infer the concentrations of individual odorants in an odor mixture. The perception of odors has been thought as synthesis, i.e., odorant mixture is perceived as a unit odor [16]. Nevertheless, the performance of the reconstruction task indicates that most of the information about the odor mixture including the identities and concentrations of individual odorants in a sparse mixture can potentially be extracted from the activity pattern of ORNs, which is consistent with the experimental finding that mice after training can detect a target odorant in odor mixtures with up to 16 different odorants [80]. In this work, we focused only on the optimal coding strategy for the peripheral ORNs. In the fly olfactory system, odorants that elicit very similar ORN response patterns can be represented by very distinct patterns of KCs [41, 60]. It remains an interesting open question whether and how the architecture of the ORN/PN to KC network optimizes the odor information transmission to enhance precision of downstream learning and decision-making.

In conventional compressed sensing theory with linear sensors, a random measurement matrix enables accurate reconstruction of sparse high dimensional input signals [37, 40]. By using prior information about the input, a better sensory matrix can be designed [81, 82]. In many cases, the optimal matrix maximizes the entropy of compressed representation [83]. Unlike the linear CS problem where the measurement matrix is known and can be used directly for reconstructing the sparse input signal by using the *L*_1_-minimization algorithm, reconstruction in the nonlinear CS problem studied here has to be done by learning without prior knowledge of the sensitivity matrix. Despite this difference, our results suggest that with nonlinear sensors, the sparse optimal sensory matrix that maximizes information transmission enables better learning and more accurate reconstruction. This general observation and the limit of reconstruction in nonlinear CS should be examined with more rigorous analysis and larger numerical simulations.

Finally, in our study, we considered the simplest case where odorants appear independently in odor mixtures. However, even in this simplest case, we have found weak but statistically relevant “orthogonal” structure in the optimal sensitivity matrix [Fig. 2(c) and Fig. S1 in SM]. In naturally occurring odor mixtures, co-occurrence of odorants in different odor sources are common. For example, odorants that are products in the same biochemical reaction pathway, i.e., fermentation, are likely to appear together [2, 84]. Although odorant-evoked ORN response patterns are not simply determined by the molecular structure, some very similar odorants do trigger similar ORN response patterns [41]. On the other hand, ORNs and their responses to different odorants can be correlated due to structural similarities in their receptor proteins. It would be interesting to explore how such correlations among ORNs and odorant molecules as well as co-occurrences among different odorants in odor mixtures can affect the optimal coding strategy at the olfactory periphery in a future study.

## ACKNOWLEDGMENTS

We thank Xiaojing Yang, Guangwei Si, Jingxiang Shen, Louis Tao, and Roger Traub for helpful discussions and comments. The work was supported by the Chinese Ministry of Science and Technology (Grant No. 2015CB910300) and the National Natural Science Foundation of China (Grant No. 91430217). The work by YT is partially supported by a NIH grant (R01-GM081747).

## Supplemental Material

### S1. ORN RESPONSE

#### A. Odor-receptor interaction and OR sensitivities

We model the odor-evoked ORN response using an equilibrium binding model, where odor receptors (ORs) have two states, ON and OFF. The activity of ORN is then determined by the fraction of ORs that are in the ON state. First, consider the case that the *i*-th receptor *R_i_* is activated by the binding of an *j*-th odorant with concentration *c_j_*, which can be described as a chemical reaction 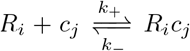, where *R_i_c_j_* is the receptor-odorant complex. At equilibrium, the activity of ORN is proportional to the fraction of odorant-bound receptor, which reads

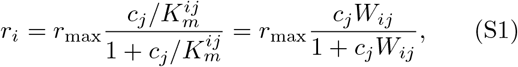

where *r*_max_ is the maximum neuronal activity, i.e., the maximum spiking rate, and we have normalized it to be 1 for simplicity. 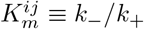 is the dissociation constant between the *i*-th receptor and the *j*-th odorant, 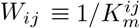 is the sensitivity constant.

Next, consider the multiple odorant molecules scenario. The response of an ORN to them can be characterized by a simple competitive binding model[1, 2], thus odorants compete for the only binding site of the receptor. The response of *R_i_* to the possible *N* odorants can be described by the following reactions:

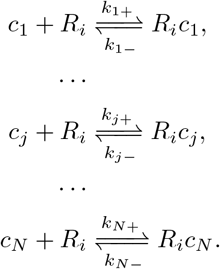

At equilibrium, the activity of *i*-th ORN is determined by the fraction of *R_i_* that are in the bound state

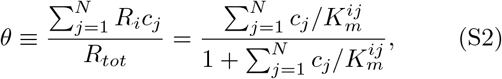

where *R_tot_* is the total number of receptors, 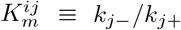 is the dissociation constant between *i*-th OR and *j*-th odorant. Hence, the response of *i*-th ORN becomes

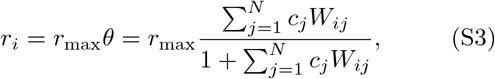

where 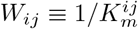.

#### B. Inhibitory response

Model for both odor-evoked excitation and inhibition have been explained in detail in [3]. Briefly, each OR has two states, ON and OFF, each of which can be further divided into two subtypes: free and odorant-bound, resulting in four different microstates. The free energy difference 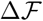 between the ON and OFF state of an OR is modulated by the odorants binding to the receptor

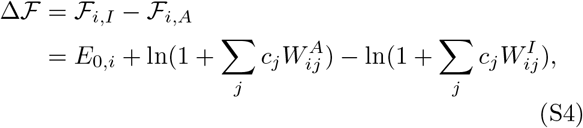

where 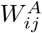 and 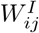 are the sensitivity of the ON and OFF state of the *i*-th OR to the *j*-th odorant. For simplicity, we have assumed that an excitatory odorant only binds to the ON state 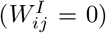, while an inhibitory odorant only binds to the OFF state 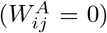. The activity of the *i*-th ORN is given by

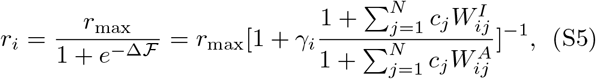

where *γ_i_* determines the basal activity 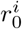 of the *i*-th OR/ORN through 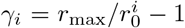, and we have omitted the Boltzmann factor *k_B_T* in the above equation for simplicity. In our model, basal activity of each receptor has been set to be the same, i.e., 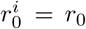, *γ_i_* = *γ*(*i* = 1,⋯, *M*).

#### C. Measured sensitivities of olfactory receptors

Responses of ORs to panels of odorants have been measured for fly larva [4], adult fly [5], mouse [6] and mosquito[7]. For fly larva, cognate odorants have been identified for each of the 18 ORs (cognate odorants could not be found for the other 3 ORs). Then a dose response curve was measured for each odorant-receptor pair. And the sensitivity of an OR to the odorant was estimated by fitting the dose response curve with a Hill function, the resulting sensitivity matrix was reported in [4]. For the mouse receptors, odor mixtures drawn from a panel of 93 odorants with various concentrations were tested against 219 receptors and 169 of them showed some level of response. A further dose-response test resulted in a response matrix for 52 ORs and 63 odorants [6]. The fraction of sensitive OR-odorant interactions were directly calculated from these sensitivity matrices.

For adult fly, the responses of 24 ORs to a panel of 110 odorants have been measured, together with the spontaneous firing rate of each ORN in the absent of stimulus [5]. Since the measurement was carried out using the same concentration of odorants, as a rough estimation, we use Eq. (S5) to fit the response and then estimate the relative sensitivity for both excitatory and inhibitory interactions. The resulting distributions of *W^A^* and *W^I^* are shown in figure 8(e), and the fraction of inhibitory interactions is about 30%. If we take a more stringent criterion and only consider the strong interactions (odorants that elicit > 50Hz response are considered excitatory odorants and odorants that suppress more than 50% of spontaneous activity are inhibitory), then the fraction of inhibition is about 11%. Different ORNs have different spontaneous firing rate, ranging from 1 spike to 52 spikes per second. The maximum spiking rate is about 294 spikes per second. Thus, the basal activity *r*_0_ estimated is in the range of 0.1 ~ 0.18, as shown in the shaded region in Fig. 8(d).

For the mosquito, the responses of 50 ORs to a panel of 110 odorants have been measured using similar method as in adult fly [7]. Since all the odorants were presented as the same concentration, we can use Eq. (S5) to fit the response and then estimate the relative sensitivities.

### S2. MEAN FIELD THEORY

In this section, we describe our mean filed theory that helps understanding the origin of the sparsity of optimal odorant-receptor sensitivity matrix in detail. We first simplify the expression of mutual information in the limit of small Gaussian noise and derive an effective target function Δ*H*. We then consider the case with only one odorant which has a broad concentration distribution and many receptors (*N* = *n* = 1, *M* ≫ 1), and next the two odorants scenario (*N* = *n* = 2, *M* ≫ 1), where we demonstrate the existence of “optimal” sparsity (*ρ_w_*) of sensitivity matrix that enables optimal coding. Finally, we extend the result to more general scenarios, i.e., *N,M* ≫ 1,*n* = 2, where the upper bound of *ρ_w_* can be approximated with the result from the case with *N* = *n* = 2,*M* ≫ 1.

As shown in Fig. 1(a) in the main text, the “coding” problem is basically how the odor vector *c* is best mapped to the ORN activity vector *r* by

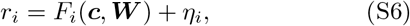

where *F_i_*(*c, **W***) is the nonlinear response function for the *i*-th neuron (*i* = 1, 2,…, *M*) as introduced in previous section, and *η_i_* is the noise, which may assume certain specific form given its biophysical/biochemical origin. For convenience, here we consider it to be a small Gaussian noise with zero mean and a standard deviation *σ*_0_, and noise among ORNs are independent.

For a given odor environment, characterized by the mixture distribution *P*_env_(***c***), we can derive the distribution of ORNs response

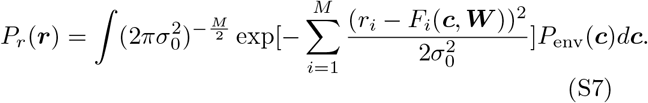

The optimal entropy coding strategy is to determine the response functions, which are characterized by the sensitivity matrix ***W***, to maximize the mutual information *I*(***c, r***) [Eq. (2) in main text] between odor environment *P*_env_(***c***) and ORN response pattern *P_r_*(***r***). In the limit of small noise, the conditional entropy *H*(***r|c***) → 0, hence *I*(***c, r***) ≈ *H*(*r*), the differential entropy in the response space is

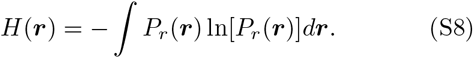

We first consider the log-normal distribution of odorant concentration, then *H*(***r***) can be simplified as

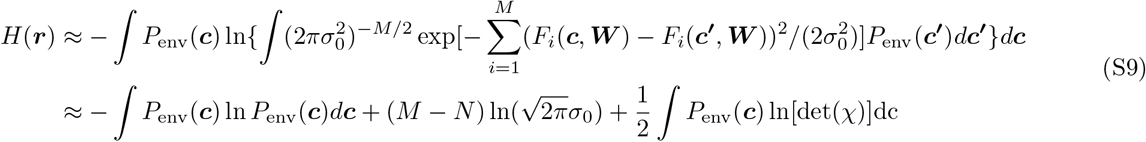

where *χ* is a *N × N* Jacobian matrix with its matrix elements given by

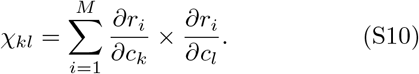

Eq.(S9) holds when *N* ≤ *M*. The first two terms in the expression for *H*(***r***) are independent of ***W*** with clear physical meanings. The first term is just the entropy of the odor distribution, and the second term is due to dimension reduction. The third term is the most interesting term. It depends on the sensitivity matrix and it is this term that we need to optimize given the odor distribution *P*_env_(***c***) and the form of the response function *F*. We denote the third term as

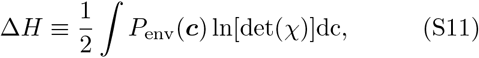

which plays the central rule in the following analysis. Here and after, we will call Δ*H* as our simplified target function which is to be maximized.

#### A. One odorant and multiple receptors

Since we are considering the log-normally distributed odorant concentration, it is convenient to work in log scale. We now define 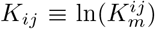 and 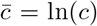. For the case *N* = *n* = 1 and *M* ≫ 1, *χ* becomes a scale, which is

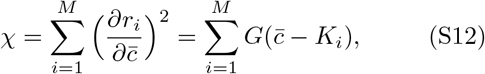

where we have defined *G*(*x*) = *e*^2*x*^/(1+*e^x^*)^4^, the square of the derivative of the response function. It has a peak at *x* = 0 with a width ~ 1, i.e., it can be well approximated as the product of the standard Gaussian function and a constant: 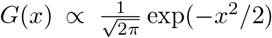. Eq. (S11) then becomes

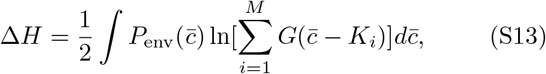

In the limit of *M* → ∞, the problem can be solved by assuming a distribution of *K, P_k_*(*K*), which leads to:

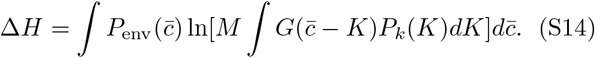

Now the problem becomes optimizing Δ*H* with respect to the distribution *P_k_*(*K*) under the constraint ∫*P_k_*(*K*)*dK* = 1. Using variational method, the problem becomes an optimization of a functional 𝓛[*P_k_*] with Lagrangian multiplier

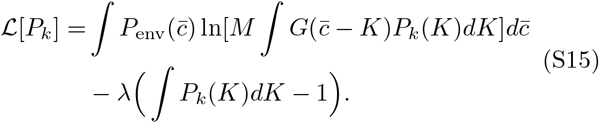

The necessary condition for the maximum value of 𝓛 with respect to *P_k_*(*K*) is

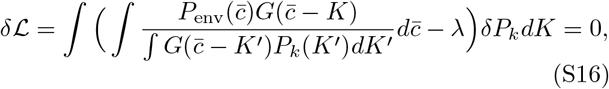

which holds for arbitrary variation *δP_k_*. So, we have

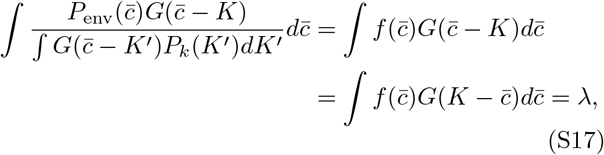

where we have defined

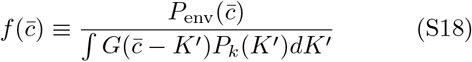

and used the fact that *G*(*x*) is an even function. To find the condition of *P_k_*(*K*) that satisfies Eq.S17, take the Fourier transform of Eq.S17, we have

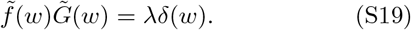

From the reverse Fourier transformation, we have

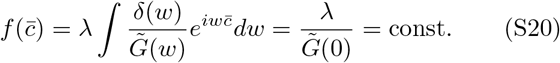

From the definition of 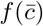 (Eq. S18), we derive the necessary condition to achieve maximum 𝓛

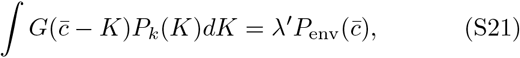

where 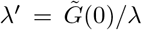 is a constant, which can be determined by integrating both sides of the above equation. Substitute 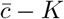 with *x* and integrate over *x*, we have

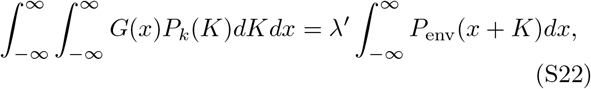

the LHS of the above equation is 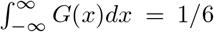, while the integration in the RHS is independent of *K* (recall that 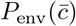 is a Gaussian function with respect to 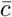) and is just 1, thus *λ*′ = 1/6.

Eq. (S21) can be solved by the deconvolution method. We first approximate 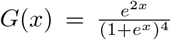 by the Gaussian function, i.e., 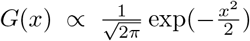(they have nearly identical shape). From the Fourier transformation of Eq. (S21), we have

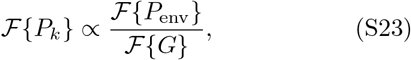

where

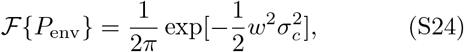

and

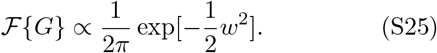

Thus

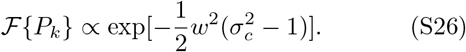

By the inverse Fourier transformation of the above equation, we have

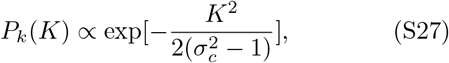

which is a Gaussian distribution with a standard deviation 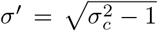. By definition, *W_ij_* = exp(−*K_ij_*), thus *W_ij_* follows a log-normal distribution, which is Eq. (3) in the main text.

As *M* → ∞, plug Eq. (S27) in Eq. (S14), we derive Eq. (4) in the main text, thus

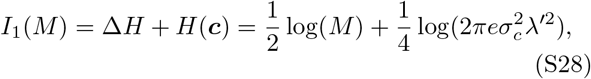

where *σ_c_* is the standard deviation of the input and the only term in *I*_1_(*M*) that depends on *M* is 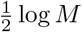. Thus, when *M* is sufficiently large, *I*_1_(*M*) increases logarithmically with *M*.

#### B. Two odorants and multiple receptors

In this scenario, we use both explicit computation and phenomenological model to approximate Eq. (S14). In order to compute Eq. (S14) explicitly, we first write down the matrix elements of *χ*

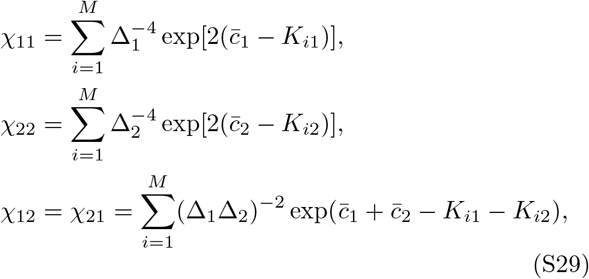

where 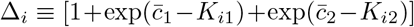 for *i* = 1, 2. From the *χ* matrix, we can calculate its determinant 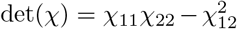 and thus obtain the expression for Δ*H*: where 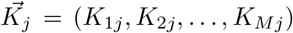 is a dimension-M sensitivity vector for *j*-th odorant in the receptor space. The optimal design can be obtained by requiring *∂ΔH/∂K_ij_* = 0 for all *j* = 1, 2 and *i* =1, 2,…, *M*.

In the limit of *M* → ∞, the problem can be tackled using a distribution function *P_k_*(*K*_1_,*K*_2_). The matrix elements of *χ* now depend on this distribution. Using variational method similar to the case for *N* = 1, the final equation for *P_k_*(*K*_1_, *K*_2_) is

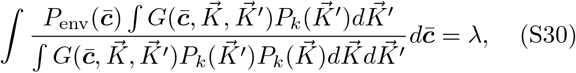

where

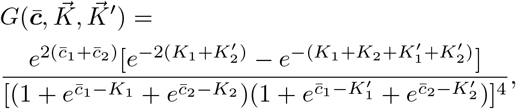

and 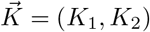 and 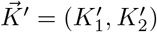 are the sensitivity vectors, whose components are the sensitivity of a given receptor to one of the two odorants. Multiply both side of Eq. (S30) with 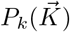 and integrate over 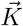, we can determine 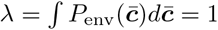.

To simplify the notation, we now define *v*_1_ = exp(−*K*_1_),*v*_2_ = exp(−*K*_2_), i.e., *v*_1_,*v*_2_ are just the sensitivity in linear scale. Now we have an explicit form of Δ*H* when *N* = 2, we shall first use appropriate approximation to demonstrate the existence of “sparseness” in sensitivity matrix ***W*** when *N* = 2, and use numerical calculations to determine how the sparsity of sensitivity matrix and the distribution of its non-zero elements depend on input statistics. Further, we extend the case to *n* = 2, *N* > 2, and demonstrate that the fraction of sensitive elements at *N* = 2 is an upper bound for scenarios where *n* = 2, *N* > 2.

For a large number of receptors and two odorants considered here, we can assume that the distributions for the two sensitivities *v*_1_ and *v*_2_ are independent with the joint distribution

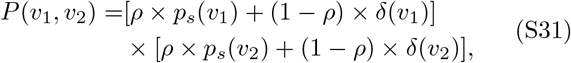

where *p_s_*(*v*) is a smooth distribution function that is the same for all odorant-receptor pairs, and *ρ* is just the fraction of non-zero elements, i.e, the “sparsity” of the matrix.

We can now write down Eq. (S29) explicitly in this particular case,

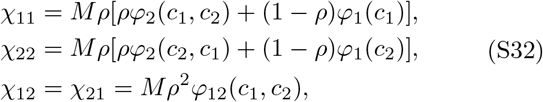

where

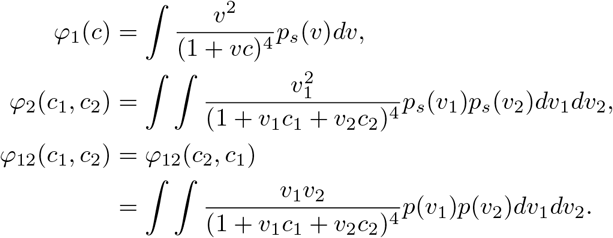

Plug Eq. (S32) into Eq. (S11), we have the expression for the differential entropy

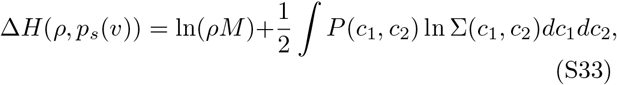

where Σ(*c*_1_,*c*_2_) is defined as:

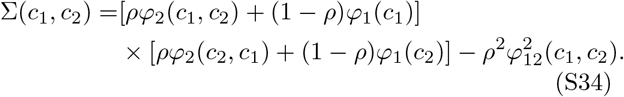

To compare with the one odor case, Eq. (S33) can be further written as

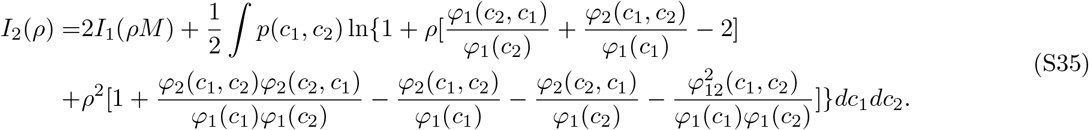

Given a value of ρ, and assuming *p_s_(v)* is the same as the one odorant case, the second term of the above equation can be numerically integrated directly. As shown in Fig. 5(b) in the main text, *I*_2_ achieves maximum value at an optimal *ρ*\* < 1.

#### C. Analytical approximations of optimal sparsity

With *N* > 2, *M* →, and *n* = 2, the solution for the optimal sensitivity matrix can be also formulated as a distribution of the weights *P*(*K*). Since the distribution of the odorants are i.i.d., it is reasonable to assume a similar symmetric structure for the optimal *P*(*K*), suggesting that *P*(*K*) should be the same for different odorants, and *P*(*K*_1_, *K*_2_) should be the same for different odorant pairs.

Since only two odorants appear in any mixtures, the leading term of the constraints on the distribution of *K* therefore comes from the “pairwise interaction”(receptors that are sensitive to both odorants are likely to be “corrupted”). For simplicity, we ignore the “interference” between different samples. As we have shall see in the next section, ignoring the “interference” term would give us an upper bound of the optimal sparsity level. Therefore we have reduced the problem to two odorants and infinite receptors. So Eq. (S33) can approximate the target function. Given the constraint ∫*p_s_(v)dv* = 1, the problem becomes optimizing the following function with respect to *ρ,p_s_(v)*:

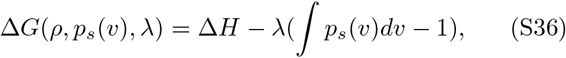

where λ is the Lagrange multiplier. Take the derivative with respect to *ρ*, we have:

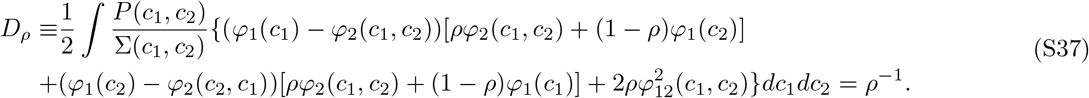

To obtain the optimal functional form of *p_s_(v)* directly through variational method is difficult, therefore we further parameterize the distribution *p_s_(v)* with a lognormal distribution, i.e., 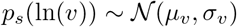, which is supported by our numerical simulation. Therefore we can take derivative of Eq. (S33) with respect to *μ_v_* and *σ_v_*

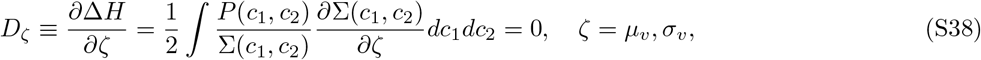

where

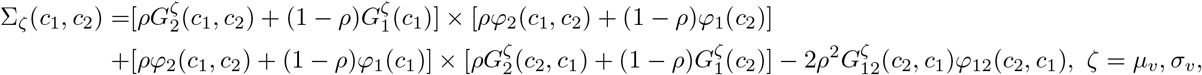

and

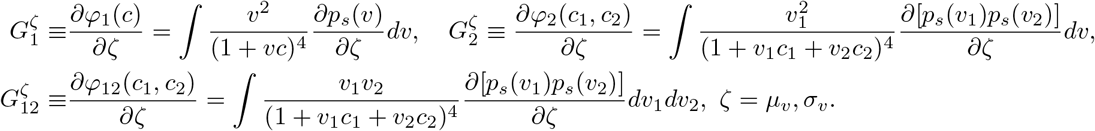

We then solve the above three coupled equations by turning the problem into an optimization problem

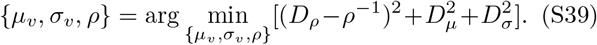

The solution is shown in Fig. S5, which agrees with the direct simulation qualitatively [see Fig. 3(a)]. The quantitative difference is due to the omission of the interference between odor samples.

#### D. The interference between mixtures of odors

Eq. (S14) can also be approximated through a phenomenological equation by considering the interference between the two odorants. As in the main text, an intuitive mean-field equation can be written down

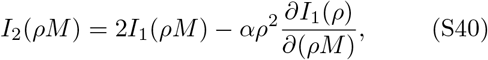

where 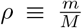 is what we defined as the “sparsity” of the sensitivity matrix with *m* the average number of receptors activated by one odorant, *α* is a phenomenological factor that quantifies how much the information conveyed by one receptor is “corrupted” when the receptor is active to two odorants in the same mixture. Comparing with Eq. (S35), we can see that *α* is a function of *ρ* and *σ_c_*, i.e., *α* = *α*(*σ_c_,ρ*). The first term of Eq. (S40) increases logarithmically with *m*, while the second term increases supralinearly with *m*, and there will be a crossover point which leads to a peak of *I*_2_ (*ρM*).

We next extend our mean-field theory to more general cases where there are *n* odorants (can be larger than 2) in odor mixtures. While unable to derive analytically how *ρ* and *p_s_* (*v*) depend on input parameters, we will show that our previous result set an upper limit of *ρ_w_*, thus introducing the interference would only cause the optimal *ρ_w_* to be an even smaller value. Starting with *N* = 3 and *n* = 2, the mutual information is

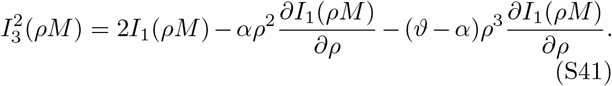

The contribution of the higher order term depend on the difference between *ϑ* and *α*, and *ϑ* is a phenomenological factor that quantifies how much the information conveyed by one receptor is “corrupted” when the receptor is active to all 3 odorants. It is obvious that *ϑ* – *α* > 0. This can be extended to more general case, i.e., *n* > 2,*N* > 2 where there are *n* odorants in each mixture and *N* possible odorants, we have

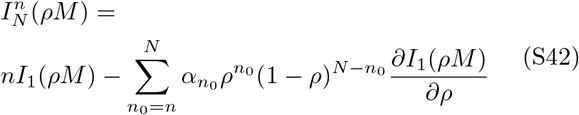

with

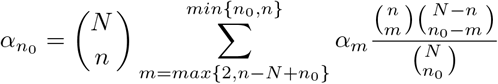

and

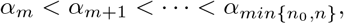

*α_m_* should increase with *m* sublinearly, because *α_m_* describes the loss of information when a receptor is active to *m* odorants, and this information has a lower bound, as *n* increases, the loss increases but saturates gradually, and therefore *α_n_0__* also increases with *n*_0_. It is easy to see from Eq. (S41) that the interference between odor mixtures adds more negative terms, thus would lead to a smaller optimal sparsity level.

### S3 NUMERICAL OPTIMIZATION PROCEDURE

In this section, we describe the procedure of numerical simulation to search for optimal sensitivity matrix in detail. We first explain how we estimate the target function, i.e., differential entropy, at each step of the iteration, and how the genetic algorithm works to find a local minimum of the target function in each trial. Then, we explain the difference between optimization procedures for the excitation-only case and for receptors with both excitatory and inhibitory response.

#### A. Estimation of differential entropy

The response of ORNs *r* is generated through Eq. (S3) using 50000 odor stimuli sampled from *P_env_*(*c*). The differential entropy of *P_r_*(*r*) is estimated using a copula-based method, the Gaussian Copula Mutual Information(GCMI) estimator[8, 9], which uses the concept of statistical copula to give a Gaussian parametric estimation of mutual information for variables with any marginal distributions. The differential entropy *I*(*r*) is the sum of the entropy of each marginal distribution *I*(*r_i_*) subtracting the joint mutual information of the random variable MI(*r*), thus

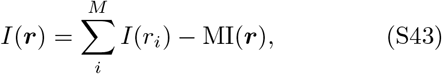

where *I*(*r_i_*) is estimated by KDP algorithm provided in the *Information theoretical estimators toolbox* [10].

#### B. Genetic algorithm for optimization

We use the covariance matrix adaptation-evolution strategy (CMA-ES) algorithm to do the numerical optimization[11, 12]. First, a population of initial candidates in the solution space are randomly sampled from a probability model, their performance is estimated according to Eq. (S43), and part of the best “performers” are selected and used to update the parameters in the probability model. Then, a new population of solutions are generated according to the updated probability model. The iteration keeps going until the solution converges. Since this method does not guarantee global optimal solution, for each combination of different parameters *N,M,n* and *σ_c_*, we perform many simulations from random starting points, and the results are robust with different initial conditions (very similar values of final target function). In all our simulations, the algorithm runs for 5000 iterations to ensure convergence.

#### C. CMA-ES algorithm

In the case where both odor-evoked excitation and inhibition exist, the sensitivity matrix ***W*** can be divided into two parts, *W^A^* and *W*^1^. Since the target function *H(r)* does not continuously change as the values of *W_ij_* go from positive to negative values, instead of directly searching in the ***W***-space, we search solutions in the “normalized response space”(response to unit concentration). Thus, the elements of *W_ij_* is mapped to a quantity *f_ij_* which we refer as a “standard response”, based on whether it is excitatory or inhibitory,

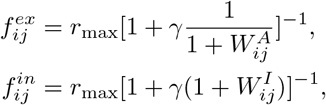

where *γ* is the constant that determines the basal activity of ORN. In this way, at each iteration, we update *f_ij_* continuously and *W_ij_* can be derived from *f_ij_* at the final stage.

### S4. DOWNSTREAM DECODING TASKS

The potential advantage of having maximum entropy coding at ORNs are demonstrated using two tasks: reconstruction and classification. For a set of parameters (*N, M, n, σ_c_*), we have found the optimal sensitivity matrix ***W***\*through numerical search. The coding scheme is changed by only changing the sparsity *ρ_w_* of ***W*** without changing the distribution of sensitive elements, i.e., they are from the same distribution as in ***W***\*. By doing so, we can control the coding capacity simple by varying *ρ_w_* and examine its consequences on downstream decoding tasks.

#### A. Reconstruction task

We use a multilayer artificial neural network (ANN) as the decoder to reconstruct odor vectors. During the training phase, 10^6^ sparse odor stimuli are generated from *P*_env_(*c*), and the activity patterns of ORNs they elicit are calculated by Eq. 1 in the main text, which are the inputs of the first layer of the decoding network. The decoder infers both the composition and intensity of odorants, so the cost function consists of two parts, the “identity error” and the “intensity error” [Eq. (7) in main text]. After training, the performance of ANN is tested using a new set of odors that are drawn from the same distribution as used in the training phase.

#### B. Classification task

Odors are presented as clusters whose centroids are sampled from *P*_env_(*c*), members in a cluster represent variations of the centroid. Denote the stimulus of a centroid as 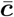, variations are generated according to 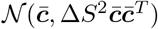, where Δ*S* is termed “radius” of a odor cluster. In each simulation, each of the 100 centroids (clusters) is randomly assigned with a category (label), i.e., attraction or aversion. Each cluster contains 50 variations with the same label as the corresponding centroid, resulting in an ensemble of 5000 stimuli, which are randomly divided into training set (80%) and testing set (20%). The representations at ORNs are used as the input of the single hidden layer decoder, 500 neurons in hidden layer (KC) and a single output neuron (MBON) are used. To model the sparse KC response observed in experiments[13, 14], numbers of connection from ORNs/PN to KC are randomly drawn from a binomial distribution, resulting on average 7 random inputs from ORNs/PNs to a KC (if *M* < 7, the maximum number of connection is *M*). The strength of connections are in the range [0,1], which are randomly drawn from the truncated Gaussian distribution 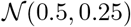. To further encapsulate the global inhibition from APL to KCs[15], following[16, 17], the input of *i^th^* KC evoked by an ORNs/PNs response vector 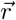 (see Eq. 1 in main text) is 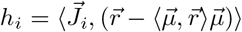, where 〈.,.〉 is the inner-product, 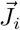 is the weight vector from ORNs/PN to KC, and 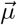 is the average ORNs/PNs response vector over all odors. The activity of each KCs is a rectified linear function of its input: Θ(*h_i_ = θ*), where Θ(*x*) = *x* if *x* > 0, and Θ(*x*) = 0 otherwise. The threshold is chosen such that on average 10% KCs respond to each odor, matching the *in vivo* experimental data[13, 18].

---

**FIG. S1.**
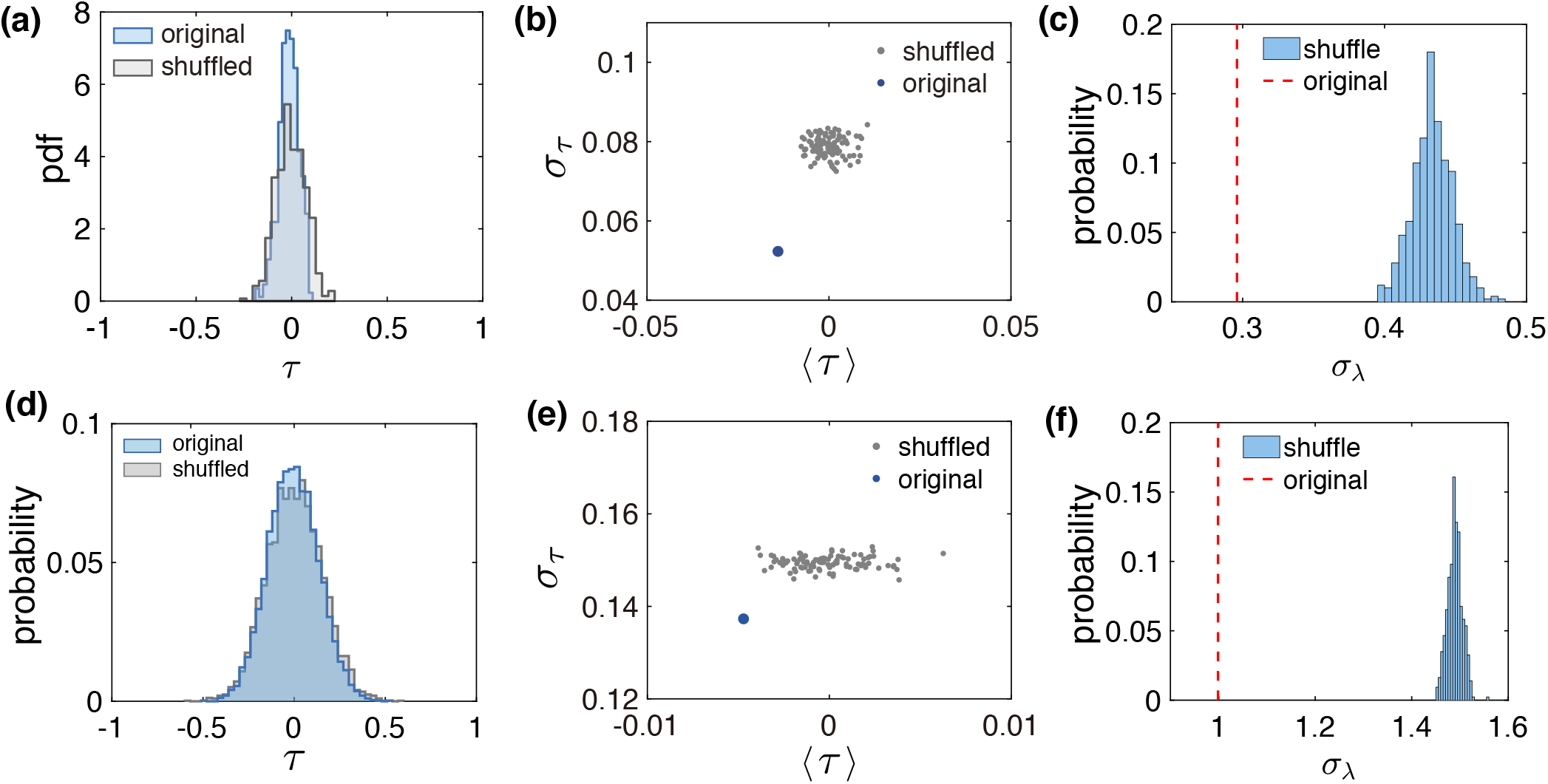
(a) Histogram of row-wise (odor-wise) Kendall’s tau rank correlation coefficient (*τ*) of the optimal matrix (blue, the matrix in Fig. 2 in main text) and its randomly shuffled matrix (gray). Note that the “insensitive” elements in the matrix has been set to be 0 before calculating the rank correlation. (b) Comparison of the average and standard deviation of *τ* for the original optimal sensitivity matrix (blue dot) and 100 randomly shuffled matrices (gray dots). (c) Optimal sensitivity matrix is more orthogonal than random matrix. Comparison of the eigenvalues of row-wise Kendall’s tau rank correlation matrix. Histogram is the standard deviation of eigenvalues from 500 randomly shuffled matrices, red dashed line corresponds to the optimal sensitivity matrix. Notice that, as a property of correlation matrix, the average of eigenvalues is 1 for both shuffled and original correlation matrix. (d) (f) correspond to (a) (c) respectively, but for column-wise rank correlations. Parameters: *N* = 100, *M* = 30, *n* = 2, *σ_c_* = 2.

**FIG. S2.**
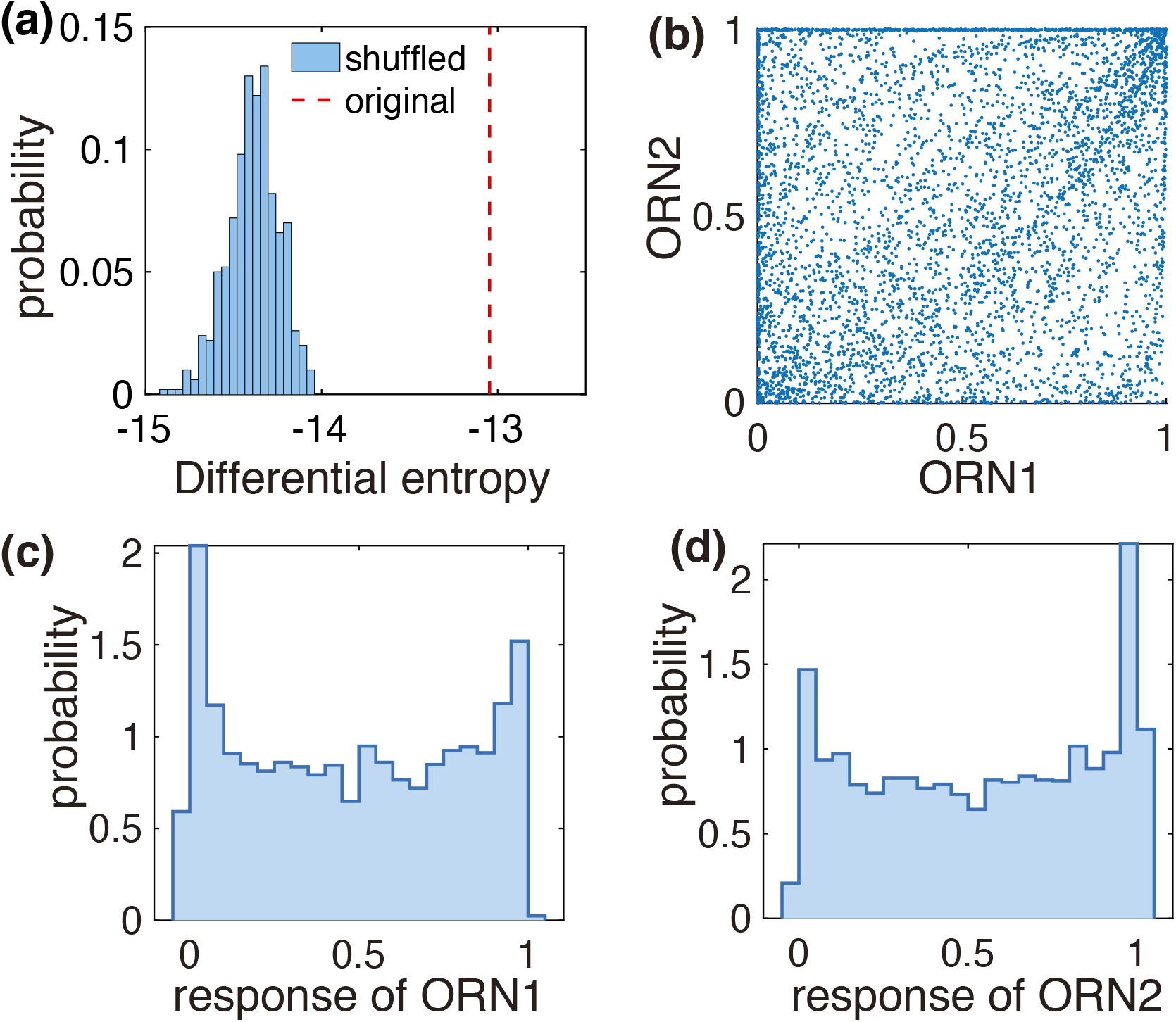
(a) Comparison of the differential entropy of ORN response I(r) with optimal sensitivity matrix (dashed line) and with randomly shuffled matrix (histogram). Parameters: *N* = 100, *M* = 30, *n* = 2, *σ_c_* = 2. (b)-(d) Optimal receptor-odor sensitivity matrix ensures near uniformly distributed and decorrelated ORN response patterns. (b) Scatter plot of responses of two ORNs to a sample of 10^4^ odor stimuli drawn from Penv(c). (c) and (d), histogram of corresponding ORN response showing roughly uniform distribution in the whole response range. Simulation parameters used in (b)-(d) *N* = 50, *M* = 13, *n* = 3, *σ_c_* = 2.

**FIG. S3.**
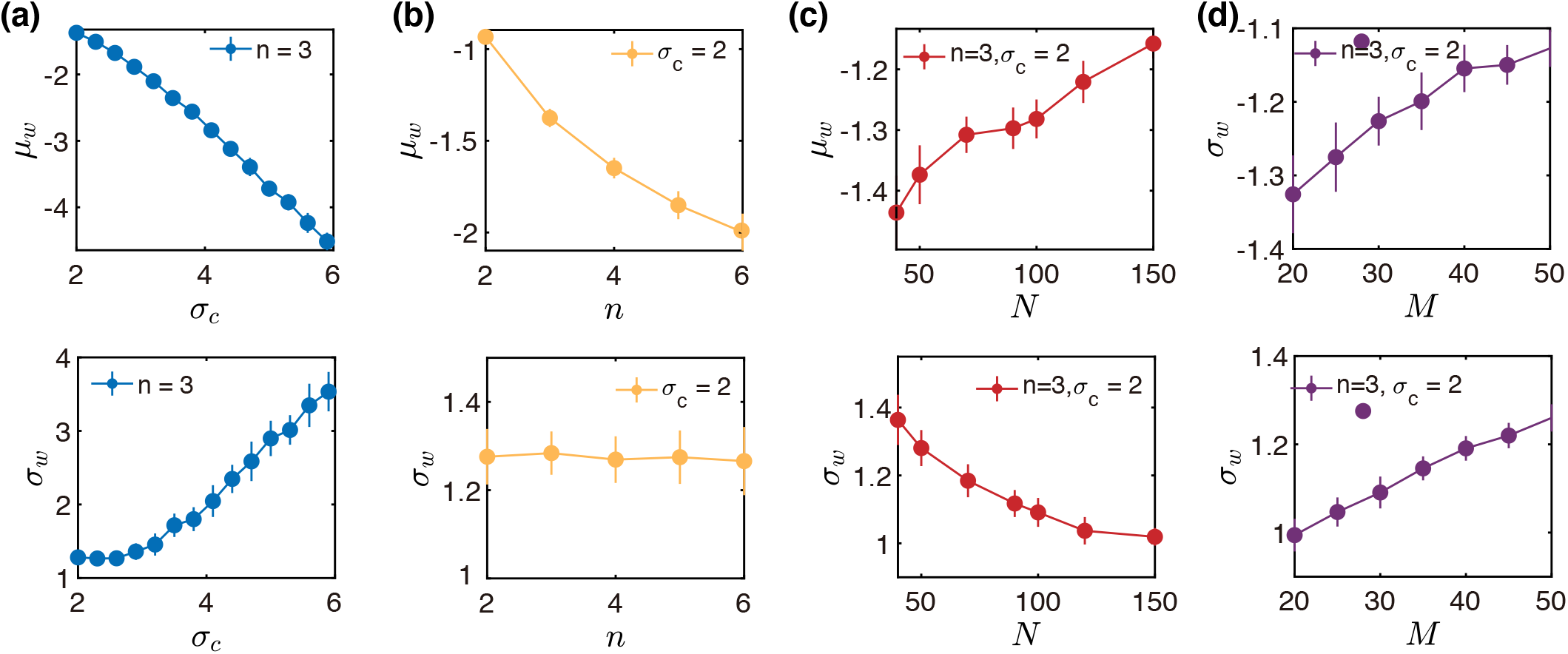
Dependence of the average *μ_w_* (upper panel) and standard deviation *σ_w_* (lower panel) of the sensitive elements in the optimal W on the width of odor log concentration *σ_c_* (a), the input sparsity n (b), the number of total odorants N (c), and the number of receptors M (d). In (a-b), N = 50, M = 13, in (c), M = 13, and in (d), N = 50. Error bars are standard deviation of 40 times simulation.

**FIG. S4.**
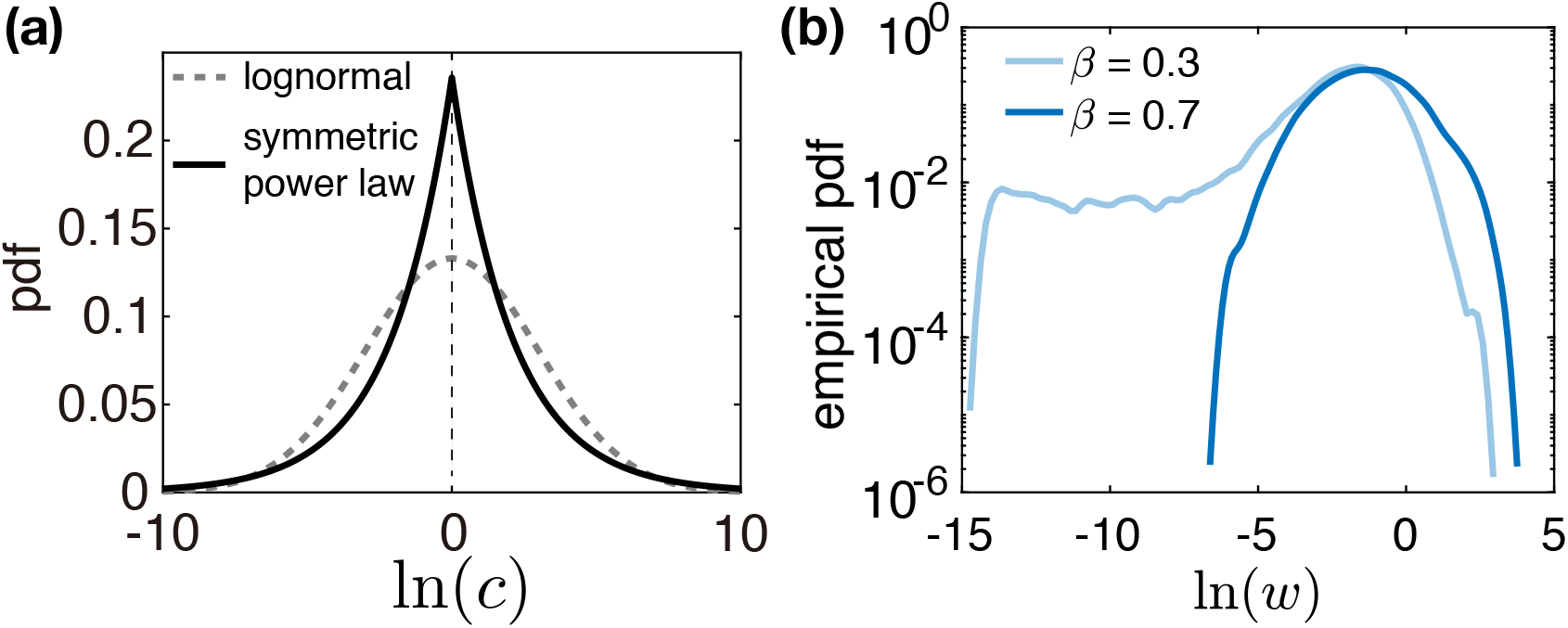
(a) Odorant concentration distribution for the symmetric power law situation (solid line), compared with log-normal distribution (dashed line) with the same variance. (b) The shape of distribution of the sensitive elements depends on the exponent of the symmetric power law. Same as figure 4 (a) in the main text, but using a log scale of the y axis.

**FIG. S5.**
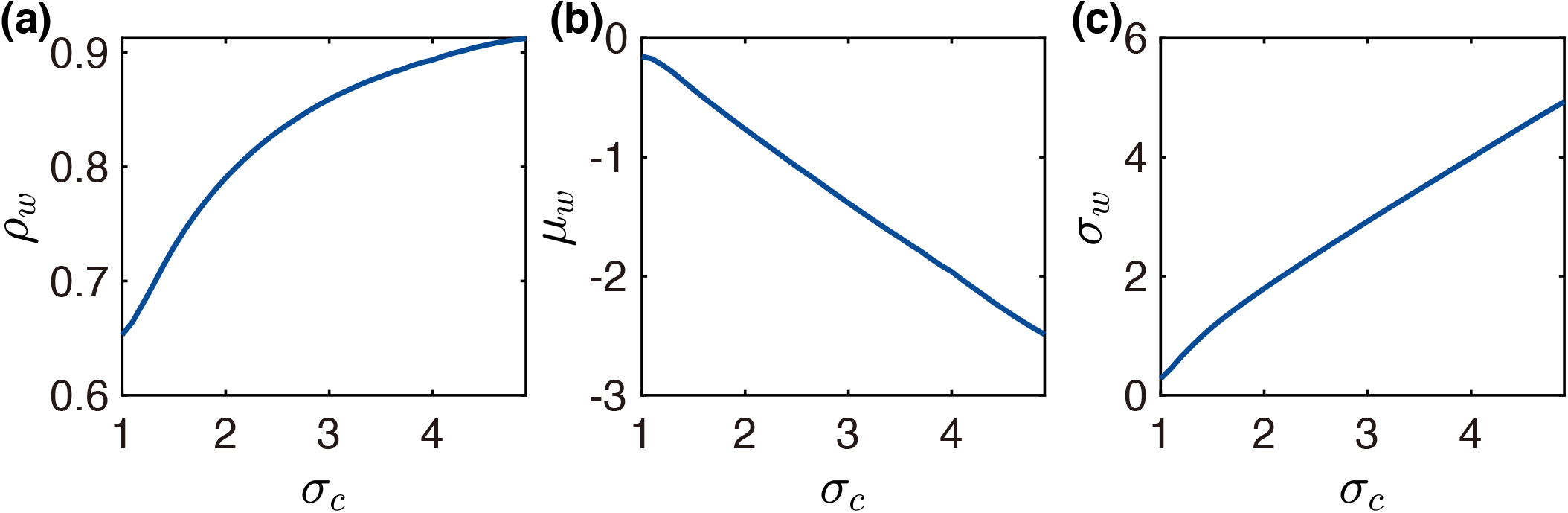
Statistics of the optimal sensitivity matrix from the mean-field theory with two odorants in mixtures. The sparsity (a), average (b) and standard deviation (c) of optimal *W* (log scale) versus the standard deviation of log-concentration of odorant. Note the qualitatively similar trends as in Fig. 3a.

**FIG. S6.**
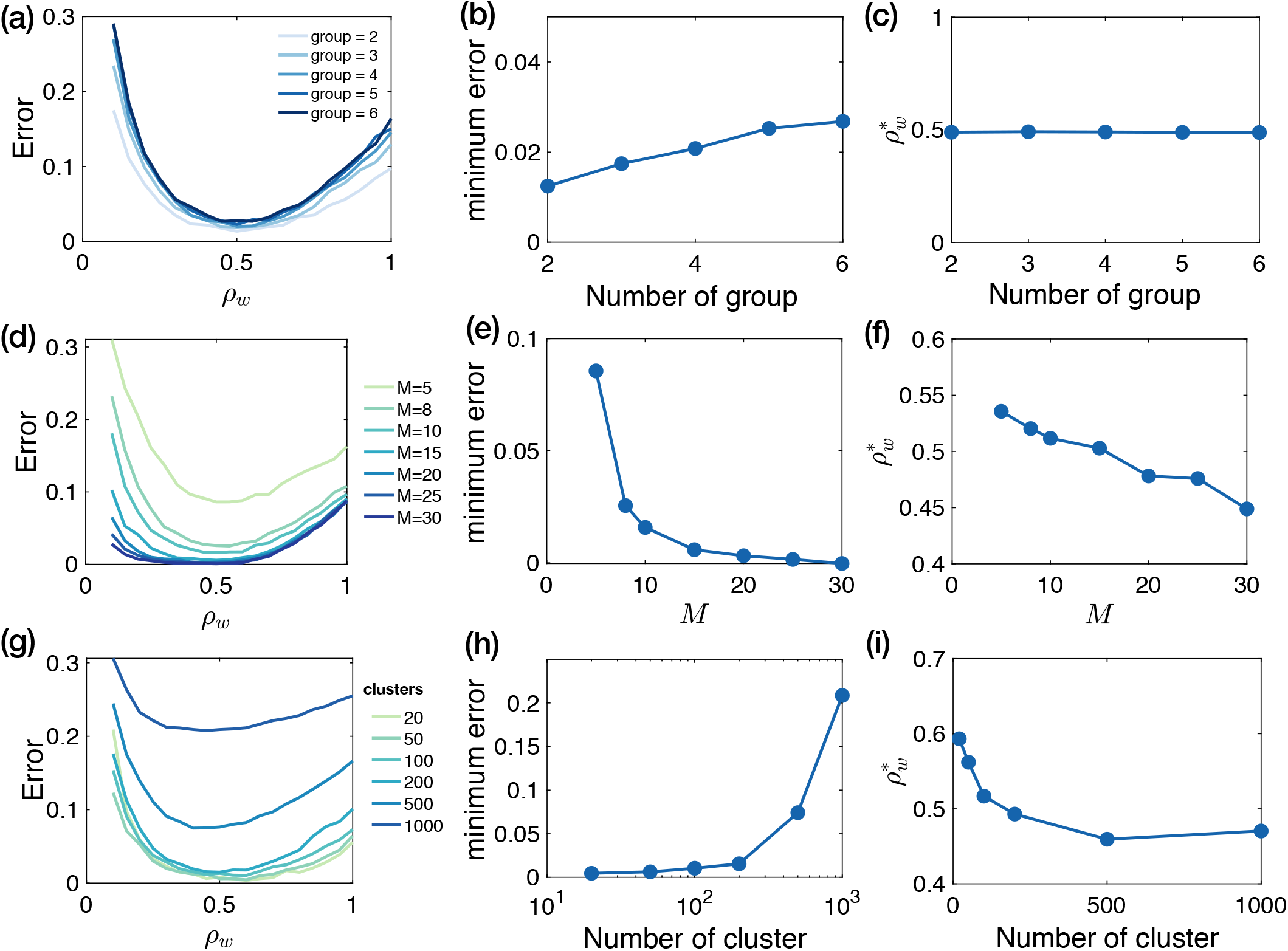
Classification performance versus the sparsity of the receptor-odor sensitivity matrix with various parameters of the decoder. (a)-(c) Classification performance for different number of groups (labels) of odor stimuli. (a) Classification error (fraction of mis-classification) v.s sparsity of W. Colors indicate different number of groups that odor stimuli have been assigned. (b) Minimum classification error with respect to number odor groups. (c) For different number of odor groups, optimal performance is achieved around *ρ_w_* = 0.5. Parameters used: *N* = 100, *M* = 12, *n* = 2, *σ_c_* = 2, Δ*S* = 0.1, *σ*_0_ = 0.1, and 100 clusters. (d)-(f) Classification performance for different number of ORNs, *M*. (d) Classification error v.s number of ORNs. Colors indicate different *M*. (e) Minimum classification error with respect to number odor ORNs. (f) For different numbers of ORN, optimal performance is achieve around 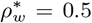 and slightly decreases as *M* increases. Parameters used: *N* = 100, *M* = 12, *n* = 3, *σ_c_* = 2, Δ*S* = 0.1, *σ*_0_ = 0.1 and two groups of labels. (g)-(i) Classification performance for different number of clusters of odor stimuli. (g) Classification error v.s sparsity of *W*. Colors indicate different number of clusters. (h) Minimum classification error with respect to number of clusters. (i) Optimal sparsity of *W* v.s the number of clusters. Parameters used: *N* = 100, *M* = 20, *n* = 3,*σ_c_* = 2, Δ*S* = 0.1, *σ*_0_ = 0.1, 100 clusters, and two groups of labels. In all these figures, line are the average classification error, error bars have been omitted to avoid clutter.

**FIG. S7.**
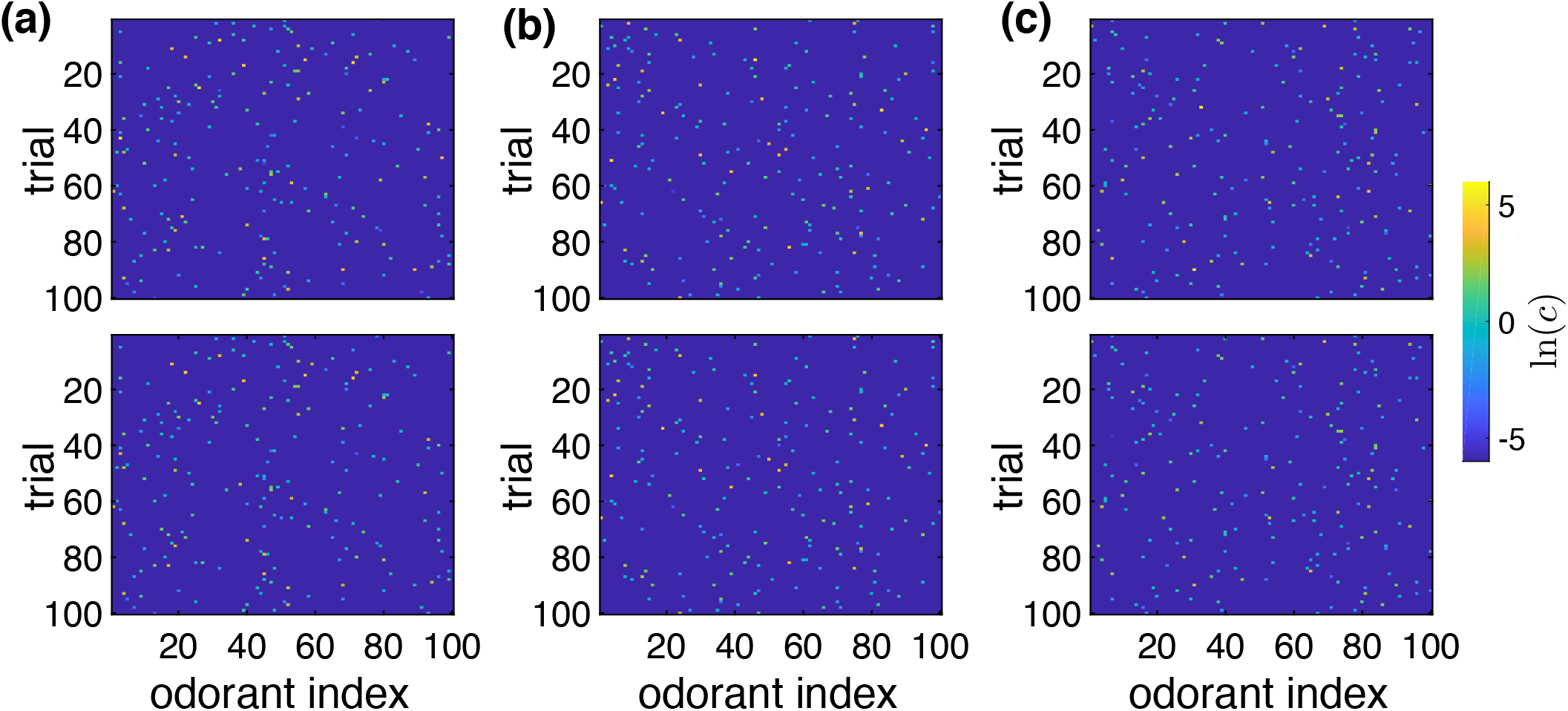
Related to Fig. 5 in main text. Comparison of the reconstructed and the input odor vector when the mapping from odor space to ORN space only differ in the sparsity of sensitivity matrix *W* (with fixed average and standard deviation of nonzero entries, *μ_w_* = −1, *σ_w_* = 2). Odor stimuli were randomly generated from P_env_(*c*). Up: 100 example sparse vectors *c* whose nonzero entries are coded with color (log scale). Bottom: the reconstructed sparse vector 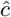. From (a) to (c), *ρ_w_* = 0.1, 0.6, 0.95 respectively. Parameters: *N* = 100, *M* = 20, *n* = 2, *σ_c_* = 2, *σ*_0_ = 0.05.

**FIG. S8.**
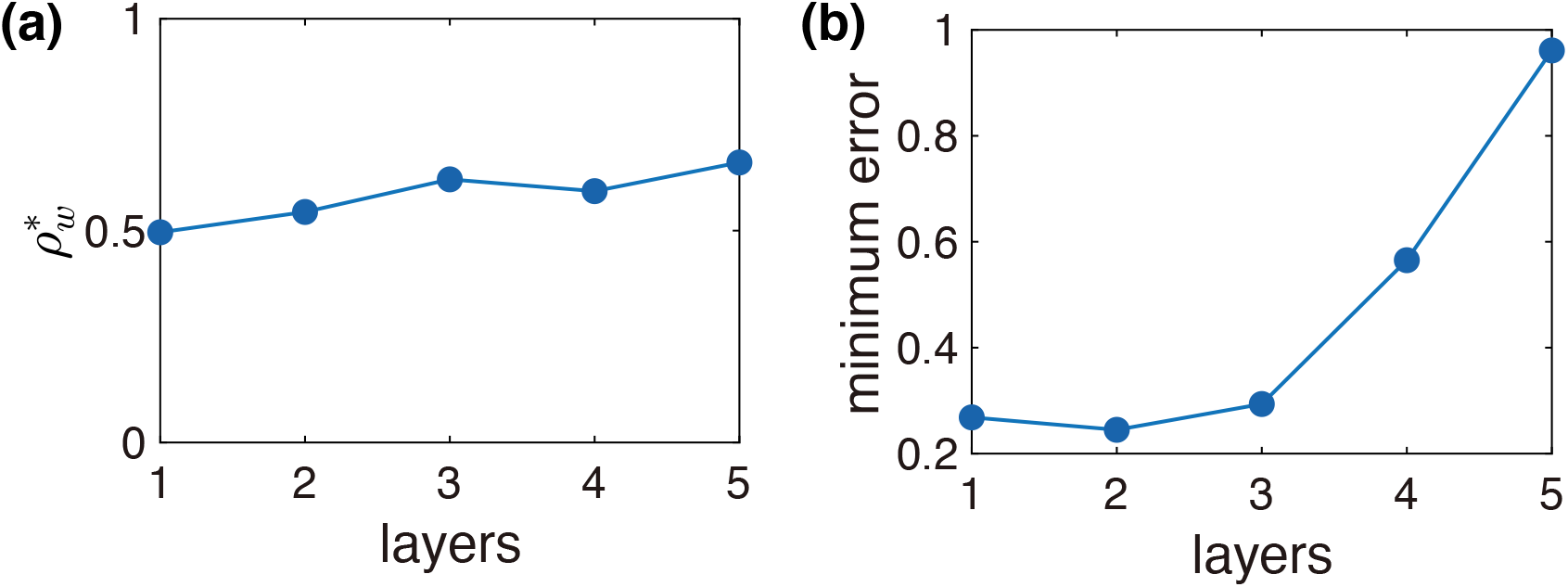
Performance of reconstruction task with different number of hidden layers. (a) Sparsity of sensitivity matrix that achieves best performance v.s the number of hidden layers in the reconstruction network. (b) Minimum reconstruction error with respect to the number of hidden layers. Parameters: *N* = 100, *M* = 20, *n* = 2, *σ_c_* = 2, *σ*_0_ = 0.05 and each hidden layer contains 100 units.

**FIG. S9.**
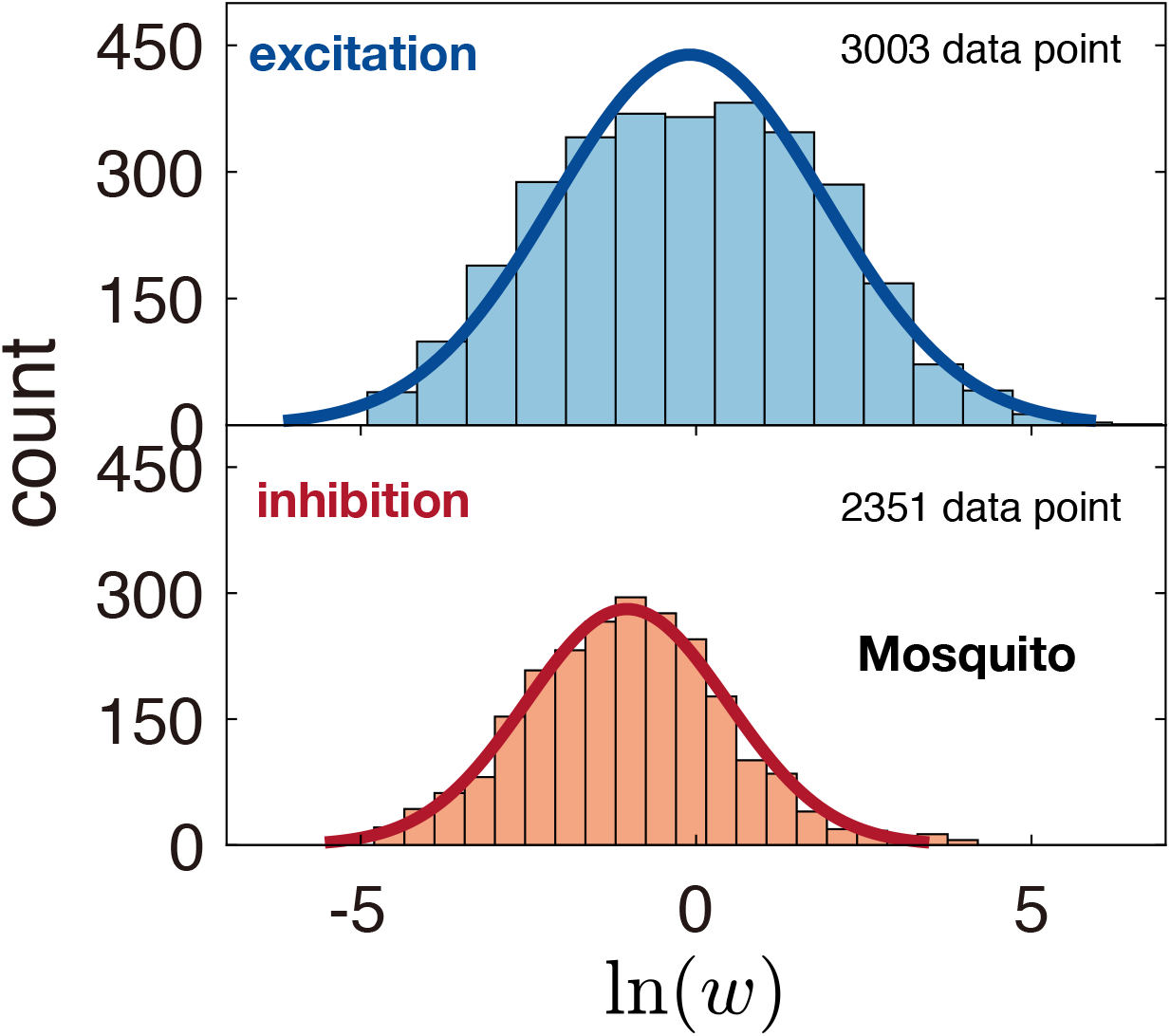
The distribution of estimated relative excitatory (upper) and inhibitory (lower) receptor-odor sensitivities from experimental data for mosquito. Solid lines are log-normal distribution fit. Data are from [7].

